# Vaccination targeting the novel endogenous tumor-associated antigen Gpr149 induces protective immunity against checkpoint inhibitor-refractory SB28 glioblastoma

**DOI:** 10.64898/2026.10.29.760792

**Authors:** Emily Steffke, Taijun Hana, Laila Latifi, Sara Abdel Malak, James McAuliffe, Vinnycius Pereira-Almeida, Nicole Briceno, Meili Zhang, Dionne Davis, Wei Zhang, Hua Song, Michael Bryan, Hye Kim, Mark Gilbert, Hideho Okada, Carol Leung, Benoit Van Den Eynde, Masaki Terabe

**Affiliations:** Laboratory of Integrative Cancer Immunology, Center for Cancer Research, National Cancer Institute, National Institutes of Health; Bethesda, MD, 20892, USA; Ludwig Institute for Cancer Research, Nuffield Department of Medicine, University of Oxford; Oxford, OX3 7DQ, UK; Neuro-Oncology Branch, National Cancer Institute, Center for Cancer Research, National Cancer Institute, National Institutes of Health; Bethesda, MD, 20892, USA; Centre for Immuno-Oncology, Nuffield Department of Medicine, University of Oxford, Oxford; OX3 7DQ, UK; Department of Neurological Surgery, University of California, San Francisco; San Francisco, CA, 94143, USA; Ludwig Institute for Cancer Research, de Duve Institute/UCLouvain; Brussels, Belgium; WELBIO Department, WEL Research Institute; Brussels, Belgium

## Abstract

**Background:** Glioblastoma is a lethal brain tumor notable for its limited spontaneous induction of CD8^+^ T cell responses. Vaccination may overcome this limitation by generating tumor-specific CD8^+^ T cells capable of mediating anti-tumor immunity in the brain. However, preclinical vaccine development is constrained by a lack of immunologically stringent glioblastoma models with defined endogenous antigens. We therefore sought to identify endogenous, vaccine-targetable antigens expressed by the immune checkpoint-refractory SB28 syngeneic mouse glioblastoma model and to evaluate their targetability using a heterologous prime-boost regimen with the chimpanzee adenovirus ChAdOx1 and the modified vaccinia Ankara (MVA) viral vector-based vaccines.

**Methods:** RNA sequencing and whole-exome sequencing were used to identify candidate tumor-associated antigens (TAAs) and neoantigens, and tumor-specific expression was validated by RT-qPCR. ChAdOx1 and MVA vectors encoding candidate antigens were generated and evaluated for immunogenicity and anti-tumor efficacy. Antigen-specific CD8^+^ T cell responses and minimal epitopes were characterized by *ex vivo* restimulation, MHC-I binding assays, tetramer staining, and tumor cytotoxicity assays.

**Results:** We identified five candidate TAAs (*Gpr149, Cdh6, Fbn2, Igf2bp1,* and *Shox2)* but no suitable neoantigen targets in SB28. Vaccination induced CD8^+^ T cell responses against the candidate TAAs, and immunogenic minimal epitopes were identified for each. Among the candidates, Gpr149 elicited the strongest vaccine-induced CD8^+^ T cell responses. In mice challenged intracranially with SB28, Gpr149-targeted ChAdOx1/MVA vaccination combined with immune checkpoint blockade significantly prolonged median survival and resulted in long-term survival in 50% of mice, whereas vaccination targeting the other TAAs did not improve survival. CD8^+^ T cells recognizing the immunodominant Gpr149_128-135_ epitope (VCYNFYTM) specifically recognized and lysed wild-type SB28 cells, but not Gpr149-knockout SB28 cells, confirming endogenous processing and presentation of the targeted epitope.

**Conclusions:** Gpr149 is an endogenous SB28 tumor antigen that can be effectively targeted by ChAdOx1/MVA vaccination to generate protective anti-tumor immunity against intracranial glioblastoma. Protection was associated with a dominant CD8^+^ T cell response to the Gpr149_128-135_ epitope VCYNFYTM. These findings establish endogenously expressed Gpr149 in SB28 as a physiologically relevant model for evaluating antigen-targeted immunotherapies and provide a systematic approach for identifying endogenous vaccine targets in other preclinical models.

**Key Messages:** **What is already known on this topic** – The immune checkpoint-refractory SB28 glioblastoma model provides a stringent setting for preclinical immunotherapy studies, but endogenous tumor antigens suitable for antigen-targeted vaccination have not been defined, requiring studies of antigen-specific immunity to rely largely on transgenic foreign antigens.

**What this study adds** – We identify Gpr149 as an endogenous SB28 tumor antigen and show that ChAdOx1/MVA vaccination generates functional Gpr149-specific CD8+ T cells that confer protective immunity against intracranial tumor challenge.

**How this study might affect research, practice or policy** – Gpr149 provides an endogenous-antigen system for more physiologically relevant testing of antigen-targeted immunotherapies in immunologically cold tumors, while the antigen-discovery approach and ChAdOx1/MVA platform provide a framework for evaluating clinically relevant glioblastoma antigens in future therapeutic studies.

## Introduction

Glioblastoma is an aggressive primary brain malignancy that remains refractory to immunotherapy despite the transformative efficacy of immune-based therapies in many other cancers^1–5^. Limited T cell infiltration and weak spontaneous CD8^+^ T cell responses are major barriers to effective anti-glioblastoma immunity. Vaccination offers a strategy to overcome this deficit by expanding peripheral CD8^+^ T cells capable of recognizing tumors. Therapeutic vaccines using peptide, nucleic acid, and dendritic cell platforms have been widely tested against glioblastoma but have yet to demonstrate improved survival in randomized trials^6–9^.

However, strategies harnessing viral vectors have focused predominantly on oncolytic virotherapy^10^, while systemic viral-vectored vaccination to generate glioma-specific T cells remains underexplored. Viral vector-based vaccines have numerous advantages, including their strong immunogenicity and their potential to encode relatively large, multiplexed inserts.

Heterologous prime-boost vaccination with the simian adenovirus ChAdOx1 followed by boost with the modified vaccinia Ankara (MVA) poxvirus induces particularly high frequencies of tumor-specific CD8^+^ T cells, making it an attractive strategy for use in the glioblastoma setting^11^. While this approach has demonstrated therapeutic potential in a variety of tumor models, leading to clinical trials for prostate cancer and NSCLC^11–15^, the ability of ChAdOx1/MVA vaccination to drive effective T cell responses in the brain is currently unknown.

The poorly immunogenic C57BL/6 syngeneic murine glioblastoma model SB28 is well-suited for preclinical testing of novel immunotherapeutic approaches. Developed within the last decade using sleeping beauty transposon-mediated oncogenesis, SB28 faithfully recapitulates many aspects of human glioblastoma when implanted orthotopically, including high levels of invasiveness, low mutational burden, and poor T cell infiltration^16^ ^17^. SB28 has therefore become the model of choice for rigorous testing of novel immunotherapies, overcoming some limitations of the commonly used glioblastoma model GL261, which is also syngeneic to C57BL/6, was created in the 1940s using methylcholanthrene, and has a high mutational burden^18–20^. While the combination of anti-PD-1 and anti-CTLA-4 checkpoint inhibitor treatment has failed repeatedly in clinical trials for glioblastoma patients and similarly elicits no response against SB28, these drugs elicit a 50% cure rate in GL261-bearing mice^16^, indicating that the GL261 model is more immunologically sensitive than human disease. The limited immunosensitivity of SB28 has established a new gold standard for preclinical glioblastoma experimentation that appears to model human treatment outcomes more faithfully than GL261.

However, specific endogenous antigens expressed by SB28 that can be targeted with vaccines are currently unknown. Potential vaccine targets include non-mutated tumor-associated antigens (TAAs) with tumor-restricted or elevated expression and neoantigens generated by tumor-specific mutations. However, neither class has been systematically defined for SB28.

Investigations of antigen-specific responses have therefore been conducted in transgenic SB28 models expressing the chicken ovalbumin (OVA) antigen^17^ ^21–23^ or human antigens including gp100 and EGFRvIII^24^ ^25^. While experimentation with xenogeneic antigen-expressing tumor models is a popular approach that has led to many important insights, the strong immune responses elicited by xenogeneic antigens can lead to an overestimation of the anti-tumor immunity of a given therapy^26^. Meanwhile, because endogenous tumor antigens are often derived from self-proteins, T cells specific for these antigens are subject to the processes of tolerance, thereby selecting against high-avidity T cell clones^27^. The threshold of T cell induction stimulated by a given endogenous antigen-targeted therapy must therefore be relatively high to observe a significant benefit.

The problem of antigenic relevance extends beyond glioblastoma to other fields with a similarly heavy reliance on approximating immunological responses based on those exerted against xenogeneic antigens. As an alternative, known murine tumor antigens can be transgenically expressed on tumor lines, such as the neoantigen *Rpl18,* which is endogenously expressed on MC38^28^ ^29^, or P1A, a murine correlate of human MAGE-type antigens^12^ ^30^. However, the strong promoters typically used to mediate transgene expression drive supra-endogenous levels of expression which may facilitate more effective CD8^+^ T cell anti-tumor immunity than would be observed against endogenous antigens with heterogeneous or low expression. To enhance the translational relevance of preclinical investigations of novel antigen-targeted therapies such as cancer vaccines, it is ideal to match the immunogenicity and expression pattern of the target antigen in the model to the relevant human disease as closely as possible. Therefore, identification of endogenous antigens in more tumor models is needed for faithful modelling of anti-tumor immunity for many cancers.

Here, we sought to identify endogenous SB28 antigens that could provide physiologically relevant targets for antigen-directed immunotherapy and to determine whether ChAdOx1/MVA vaccination against these targets could generate protective immunity against intracranial SB28. We developed a pipeline integrating tumor-selective expression, MHC-I epitope prediction, vaccine immunogenicity, endogenous antigen recognition, and anti-tumor activity. This approach identified Gpr149 as a targetable endogenous SB28 antigen and enabled evaluation of ChAdOx1/MVA vaccination in a stringent, antigenically relevant glioblastoma model.

## Results

### Identification of tumor antigen candidates in the SB28 glioblastoma cell line

While SB28 has been previously shown to harbor fewer mutations with the potential to serve as antigens than GL261^16^, specific immunogenic epitopes suitable for immunotherapeutic targeting have not been identified for the SB28 cell line. We therefore developed a pipeline to identify endogenous neoantigens and TAAs in SB28 (Figure 1A).

**Figure 1.**
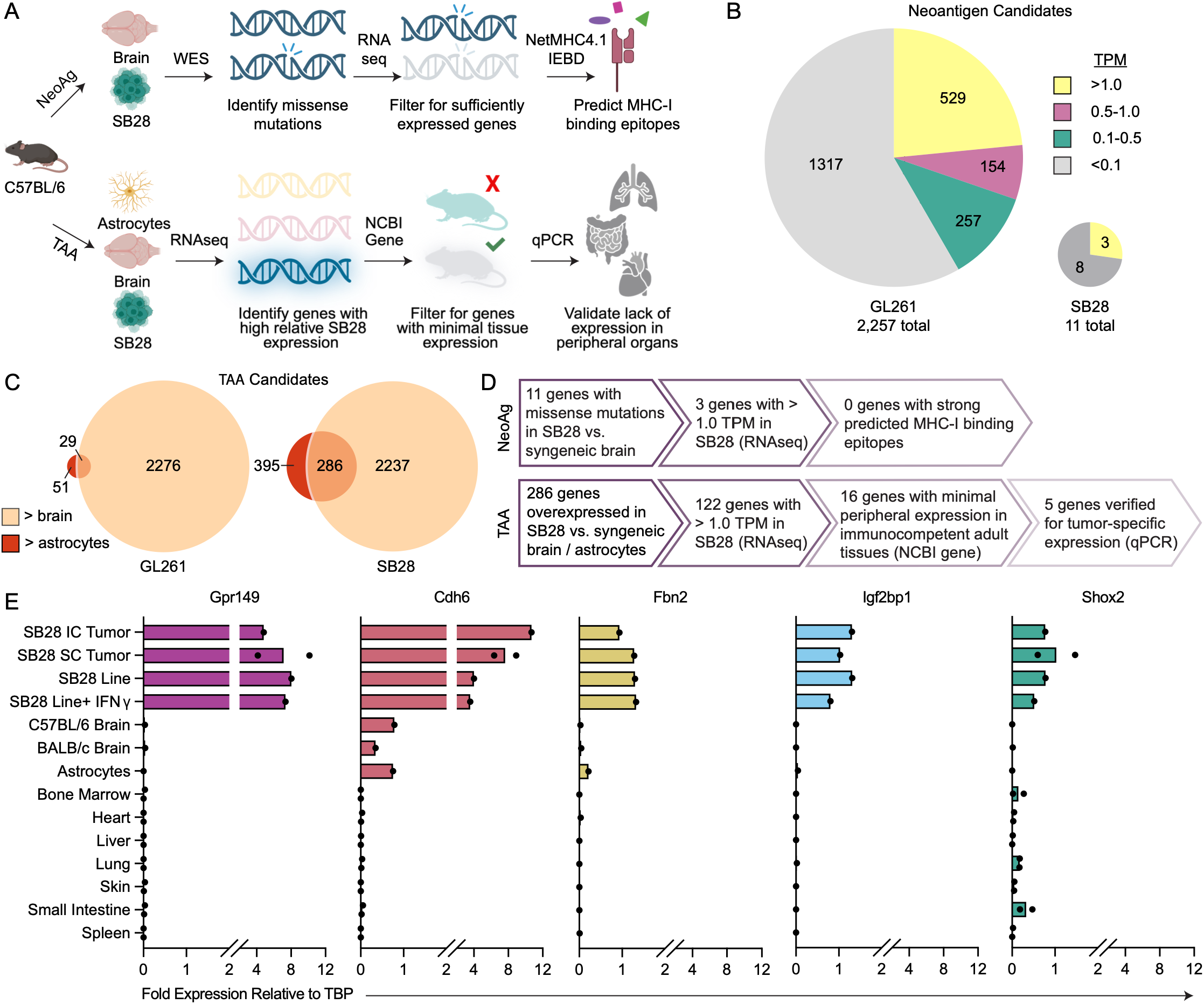
Identification of tumor antigen candidates in the SB28 glioblastoma cell line. (**A**) Schematic depicting pipeline for identification of neoantigen (*top*) and TAA (*bottom*) candidates. (**B**) Pie charts depicting the number of neoantigen candidate genes in SB28 and GL261 cell lines. Candidates were determined as missense mutations in protein-coding genes based on comparative whole exome sequencing (WES). Relative expression level of each neoantigen candidate based on RNA-seq data is summarized by color (Yellow > 1.0 Transcripts per Million (TPM), Pink 0.5-1.0 TPM, Aqua 0.1-0.5 TPM, Gray < 0.1 TPM). (**C**) Venn diagrams depicting relative numbers of overexpressed genes in SB28 and GL261 cell lines based on comparative RNA-seq, with area of circles proportional to the number of genes. Dark circles represent genes with > 0.5 TPM in tumor cells and < 0.5 TPM in astrocyte cells; light circles represent genes with > 0.5 TPM in tumor cells and < 0.5 TPM in whole C57BL/6 brain; overlap represents genes overexpressed in tumor cells compared to both astrocytes and whole brain. (**D**) Flow chart reporting numbers of neoantigen (*top*) and TAA (*bottom*) candidates passing each stage of the pipeline for SB28. (**E**) Fold change in TAA candidate gene expression in various tissues or cell lines relative to the housekeeping gene *(Tbp)* as measured by RT-qPCR. Dots represent unique tissue samples from individual mice (*n =* 1-2 / test condition). TBP, TATA-Box Binding Protein; IC, intracranial; SC, subcutaneous.

To identify neoantigen candidates, we performed whole exome sequencing and used the Mouse nEoanTigen pRedictOr (METRO) pipeline^23^ ^31^ to identify missense and frameshift mutations in SB28 or GL261 cells compared to C57BL/6 brain. Because therapeutic targeting requires sufficient antigen expression, we also performed mRNA sequencing on the same samples to evaluate the expression of neoantigen candidate genes. While GL261 cells harbored 2,257 missense or frameshift mutations, 940 of which were expressed at >0.1 transcripts per million (TPM), SB28 cells harbored over 200-fold fewer mutations, with only 11 missense and no frameshift mutations, 3 of which were expressed at >0.1 TPM (Figure 1B). Expression of all 11 SB28 neoantigen candidates is summarized in Supplemental Figure 1A.

Because CD8^+^ T cell recognition requires major histocompatibility class I (MHC-I) presentation, we submitted amino acid sequences containing each mutated residue flanked by 13 upstream and downstream amino acids to the NetMHC4.1 and IEDB Consensus algorithms to predict 8-14mer epitopes binding H-2D^b^ and H-2K^b^. Epitopes with a Consensus Percentile Rank ≤2.0 are shown in Supplemental Table 1; epitopes with an EL Rank ≤2.0 are shown in Supplemental Table 2. The genes *1700029F12Rik*, *Duoxa2*, *Fhl5*, and *Map3k19* each harbored one predicted “strong binder” with an EL Rank <0.5, and *Fhl5* and *Rabgap1l* each contained an epitope with a Consensus Percentile Rank <0.5, but none of these genes were highly expressed by RNA-seq. Among highly expressed genes, *Col1a2* and *Nras* had no predicted MHC-I binding epitopes, and *Sc5d* had one predicted epitope, HKP**Q**HIWKI, with an EL Rank of 1.462.

As genes on tumor lines may be differentially expressed in culture as compared to *in vivo*, we validated the expression of neoantigen candidates by RT-qPCR using cDNA from intracranial and subcutaneous SB28 tumors, as well as cultured SB28 cells with or without IFN-*γ* treatment. *Duoxa2*, *Fhl5*, *Map3k19,* or *Serpina1a* were not detected in the SB28 samples, and expression of *Rabgap1l* and *Fancc* was >5-fold lower than the housekeeping gene *Tbp* (TATA-box binding protein) in intracranial SB28 tumors (Supplemental Figure 1B). Among neoantigen candidates with “strong” or “weak” predicted MHC-I binding epitopes, only *Sc5d* had higher expression than *Tbp,* though expression of *Sc5d* was reduced in intracranial SB28 tumors compared with cultured cells. Overall, based on the lack of both high expression and strongly predicted MHC-I binding, no SB28 neoantigen candidates were considered promising targets.

We therefore evaluated TAAs overexpressed in SB28 relative to syngeneic brain and astrocytes. RNA-seq identified genes with a mean expression TPM >0.5 in SB28 cells and <0.5 in brain or mouse astrocytes. We identified 2,237 and 395 genes upregulated in SB28 compared with brain and astrocytes, respectively, 286 of which were shared (Figure 1C). SB28 harbored ten-fold more TAA candidates than GL261, which had 2,276 and 51 genes upregulated compared with brain and astrocytes, respectively, 29 of which were shared. To further narrow candidate SB28 TAAs, we retained genes with expression >1.0 TPM in SB28 (122 genes).

To minimize the potential for off-target toxicity, we queried the Mouse ENCODE transcriptome dataset via the NCBI Gene database and disqualified any TAA candidates with >2.5 reads per kilobase per million mapped reads (RPKM) in immunocompetent adult tissues from further consideration (Supplemental Figure 2A, B). Expression in tissues with limited MHC-I expression, including ovary, testis, and placenta, was not considered exclusionary. Of the 122 genes, 16 had minimal off-target expression and were retained for validation (Figure 1D, Supplemental Table 3). Of these, many displayed a cancer-germline-like expression pattern, with expression restricted primarily to gonadal or embryonic tissues (Supplemental Figure 2C).

To validate tumor-specific expression of these TAA candidates, we performed RT-qPCR on SB28 cells and tumor samples as well as the brain, astrocyte, bone marrow, heart, liver, lung, skin, small intestine, and spleen from healthy mice. Of the 16 candidates tested, *Gpr149, Cdh6, Fbn2, Igf2bp1,* and *Shox2* had high relative expression in SB28 and minimal off-target expression (Figure 1E). *Gpr149* showed the most tumor-selective expression, with 4- to 10-fold higher expression than *Tbp* across SB28 samples and no expression in the tested healthy tissues. Ten other genes were excluded based on low expression in intracranial tumor samples or high relative expression in off-target tissues; *IL-11* was rejected because it encodes a cytokine (Supplemental Figure 3). Of the five prioritized TAAs, *Cdh6*, *Fbn2,* and *Shox2* were also overexpressed in human TCGA glioblastoma samples (Supplemental Table 4).

### Immunogenicity of candidate tumor-associated antigens

To test the immunogenicity and anti-tumor efficacy of the TAA candidates, we created viral vector vaccines encoding the TAAs. We hypothesized that the ChAdOx1 and MVA heterologous prime-boost vaccination, which induces robust anti-tumor CD8^+^ T cell responses^11^ ^12^ ^32^, would generate high levels of SB28 TAA-specific CD8^+^ T cells.

Unlike neoantigens, which are limited to a narrow set of epitopes containing mutated amino acids, TAAs may contain a broad array of immunogenic epitopes distributed throughout the gene. To determine the presence and regionality of potential CD8^+^ epitopes in the SB28 TAA candidate genes, we analyzed the full amino acid sequence of each antigen using NetMHC4.1 and the IEDB Consensus algorithm to predict 8-11-mer epitopes binding H-2K^b^ or H-2D^b^. All TAAs contained multiple epitopes with an EL rank < 0.5 (NetMHC4.1) and/or Consensus Percentile Rank < 1.5 (IEDB) distributed throughout the sequence (Figure 2A). We therefore designed our vaccine constructs to encode the complete sequences for each TAA where feasible. Advantageously, ChAdOx1 and MVA can encode much longer gene inserts than peptide or mRNA vaccines, with a maximum insert size of approximately 5000 bp. We created four matched pairs of ChAdOx1 and MVA vaccines containing identical antigen inserts (Figure 2B). Each construct included the MHC class II-associated invariant chain transmembrane domain, which enhances CD8^+^ T cell responses against antigens encoded by viral vectors^11^ ^33^. *Gpr149* and *Cdh6* were encoded as synthetic sequences codon-optimized for mouse. As *Fbn2* exceeds 2,800 amino acids, we selected 11 minigenes centered on CD8^+^ T cell epitopes with the lowest combined prediction scores and encoded them in a codon-optimized synthetic construct separated by GGGPGGG spacers (Supplemental Table 5). The “Combi” construct included a 27-mer sequence encoding the *Sc5d* neoantigen together with full-length *Shox2* and *Igf2bp1*.

**Figure 2.**
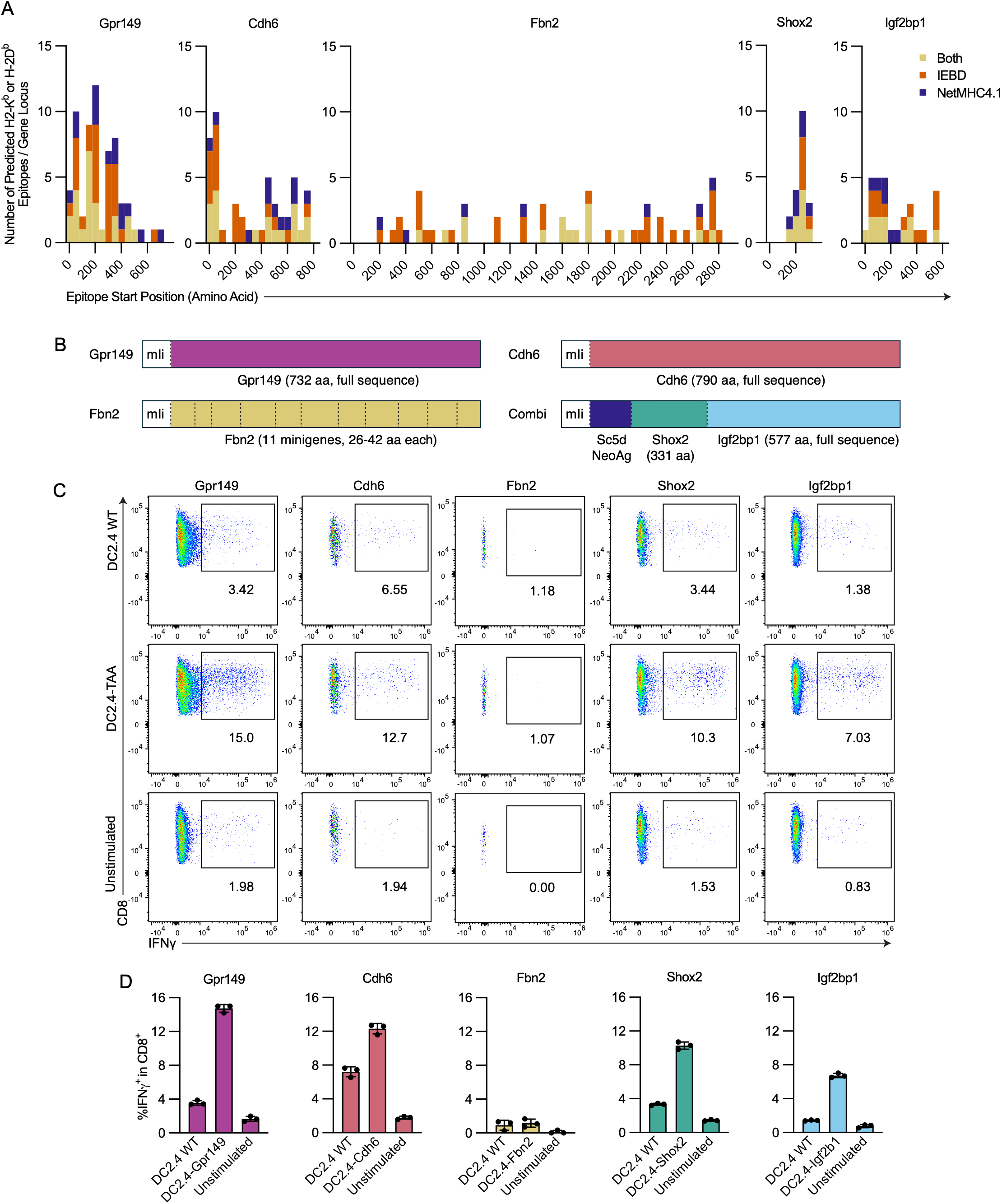
Immunogenicity of candidate tumor-associated antigens. (**A**) Histograms depicting number and distribution of unique 8-11mer epitopes beginning within a given range of amino acids (bin width = 50) for each gene predicted to bind H-2K^b^ or H-2D^b^ by the IEDB Consensus Algorithm (orange, Consensus Percentile Rank <1.5), NetMHC4.1 (navy, EL Rank < 0.5), or both (yellow). (**B**) Schematics representing the design of each vaccine. Dotted lines represent spacer amino acid sequences between gene segments (GGGPGGG). mIi, major histocompatibility complex (MHC) class II-associated invariant chain. (**C**) Representative FACS plots and (**D**) frequency of CD8^+^ T cells expressing IFN-*γ* after co-culture of *ex vivo* expanded T cells from spleens of mice vaccinated with a given ChAdOx1-TAA vaccine and restimulated with WT or transgenic DC2.4 cells expressing the cognate gene insert. FACS plots are pre-gated on Singlets/Live/CD3^+^CD8^+^. Dots in bar graphs represent co-culture replicates.

We next tested whether vaccine-encoded TAAs contained immunogenic CD8^+^ epitopes that were endogenously processed and presented. Splenocytes from mice vaccinated with two doses of ChAdOx1-Gpr149, -Cdh6, or -Combi vaccines were expanded with irradiated DC2.4 cells expressing the corresponding TAA, then restimulated with WT or TAA-expressing DC2.4 cells and analyzed for IFN-*γ* production. CD8^+^ T cells from vaccinated mice were successfully expanded by culture with DC2.4 cells expressing Gpr149, Cdh6, Shox2, and Igf2bp1, but not by those expressing Fbn2, indicating limited processing and presentation of MHC-I restricted Fbn2 epitopes. The expanded cells produced IFN*γ* in response to co-culture with corresponding TAA-expressing DC2.4 clones, demonstrating endogenous processing and presentation of immunogenic Gpr149, Cdh6, Shox2, and Igf2bp1 epitopes on H-2^b^ molecules (Figure 2C, D).

### Identification of immunogenic minimal CD8^+^ T cell epitopes within TAAs

We next identified immunogenic epitopes within the TAAs. We performed ICS on splenocytes from ChAdOx1/MVA-TAA vaccinated mice stimulated *ex vivo* with peptides corresponding to the strongest predicted MHC-I binders within each TAA (Supplemental Table 6). Peptide pools from Gpr149, Igf2bp1, or Fbn2 significantly induced IFN-*γ* in CD8^+^ T cells, with responses mapped to Gpr149_127-135_, Gpr149_128-135_, Gpr149_447-455_, Igf2bp1_83-92_, and Fbn2_1455-1463_ (Figure 3A). These peptides also induced TNF (Supplemental Figure 4A). The Shox2 pool and Shox2_145-152_ significantly induced TNF but not IFN-*γ*, and additional Gpr149, Igf2bp1, and Fbn2 peptides also induced TNF. Responses to the Cdh6 pool and Cdh6_628-636,_ Cdh6_629-636,_ and Cdh6_675-683_ were observed in some mice but did not reach statistical significance. No CD8^+^ T cell responses were detected against the *Sc5d* neoantigen, and no cytokine responses were detected in CD4^+^ T cells, which was expected given the short peptides tested (Supplemental Figure 4B). Three long Fbn2 peptides containing the top predicted H-2I-A^b^ binding epitopes were also tested (Supplemental Table 6), but none induced CD4^+^ T cell responses.

**Figure 3.**
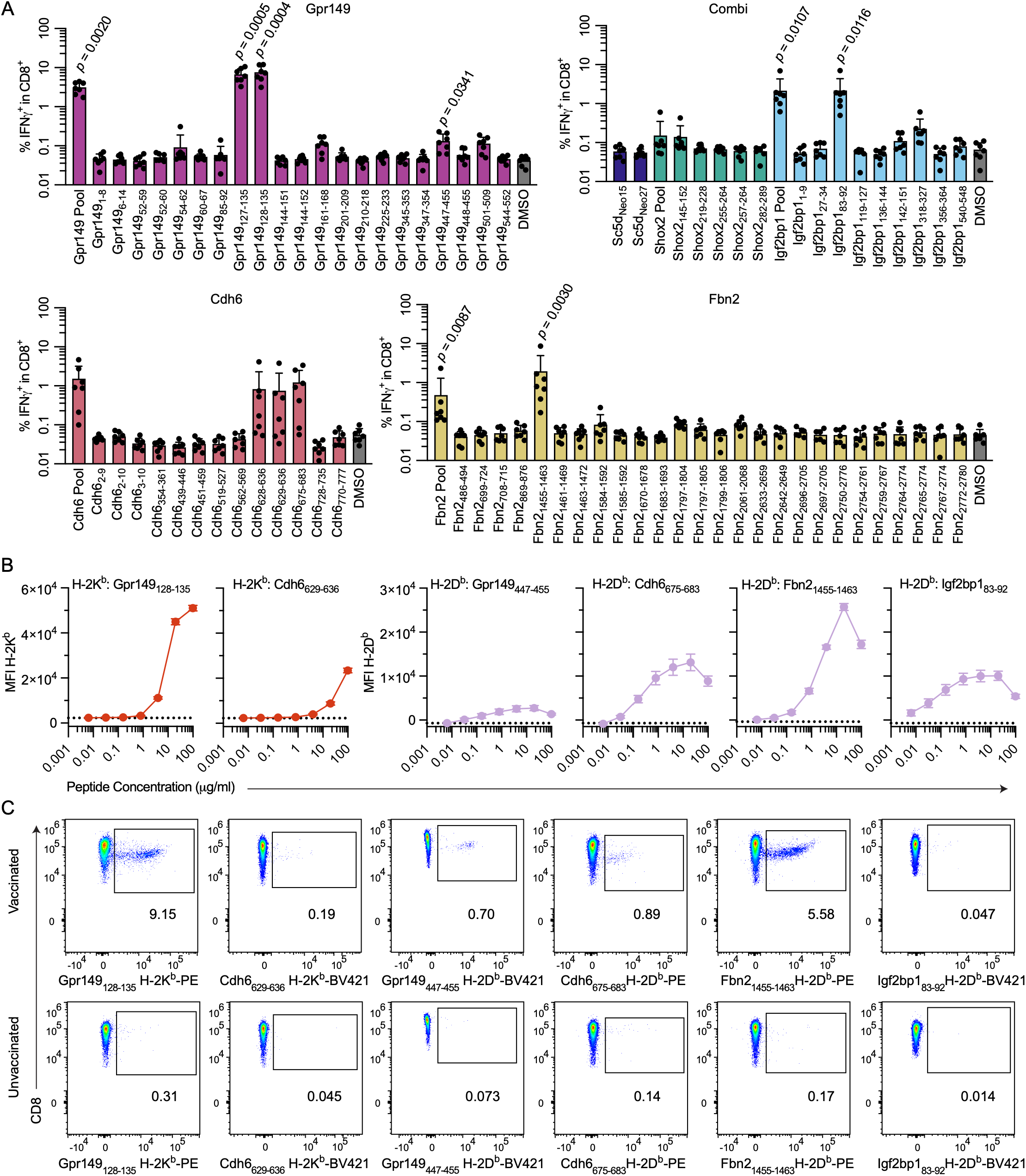
Identification of immunogenic minimal CD8^+^ epitopes within TAAs. (**A**) Frequency of CD8^+^ T cells expressing IFN-*γ* after *ex vivo* restimulation of splenocytes from vaccinated mice with minimal epitope peptides. Peptide pools include all peptides for a given gene (4μg/ml each). Dots represent individual mice (*n =* 7/group); pooled from 2 independent experiments. Statistical significance was calculated using a Kruskal-Wallis test with Dunn’s multiple comparisons for each stimulation condition vs. DMSO controls. (**B**) Median fluorescence intensity (MFI) of surface H-2D^b^ or H-2K^b^ on RMA-S cells measured by flow cytometry after overnight culture in the presence of human β2 microglobulin and peptide at increasing concentrations. Error bars represent standard deviation of triplicates, representative of 2 independent experiments. (**C**) Representative FACS plots depicting tetramer-labeled CD8^+^ T cells from blood or spleens of untreated mice or mice vaccinated with the relevant set of ChAdOx1/MVA-TAA vaccines. FACS plots are pre-gated on Singlets/Live/CD3^+^CD8^+^.

To confirm MHC-I binding and allele restriction, we loaded TAP-deficient RMA-S cells with increasing concentrations of each minimal epitope peptide and measured surface H-2K^b^ and H-2D^b^ by flow cytometry. As predicted *in silico*, Gpr149_128-135_ and Cdh6_629-636_ bound H-2K^b^, while Gpr149_447-455_, Cdh6_675-683_, Igf2bp1_83-92_, and Fbn2_1455-1463_ bound H-2D^b^ (Figure 3B), with minimal stabilization of the alternate allele (Supplemental Figure 5A). Other tested peptides failed to stabilize either allele at concentrations <20 μg/ml, consistent with low MHC-I binding affinity (Supplemental Figure 5B).

For epitopes with sufficient MHC-I binding affinities, we generated peptide-MHC tetramers and stained splenocytes or PBMCs from mice vaccinated with the corresponding ChAdOx1/MVA-TAA vaccines. Distinct tetramer^+^ populations were detected in vaccinated but not unvaccinated mice (Figure 3C). Tetramer^+^ frequencies among CD8^+^ T cells were generally similar to IFN-*γ*^+^ frequencies after *ex vivo* restimulation (Figure 3A), with the exception of Igf2bp1_83-92_, which yielded markedly fewer tetramer^+^ than IFN-*γ*^+^ cells.

### ChAdOx1/MVA-Gpr149 vaccination drives superior immunogenicity and control of intracranial SB28 tumors compared to other TAA vaccines

We then evaluated whether ChAdOx1/MVA heterologous prime-boost vaccination against SB28 TAAs could improve outcomes after intracranial SB28 challenge. C57BL/6 mice were primed intramuscularly with one of the four ChAdOx1-TAA vaccines or with ChAdOx1 expressing an irrelevant antigen (DPY), boosted intravenously 4 weeks later with the corresponding MVA vaccine, then challenged intracranially with SB28 two weeks post-boost (Figure 4A). Because constitutive expression of MHC-I on SB28 cells is minimal, the cells were pre-treated with IFN-*γ* to enhance MHC-I expression^16^. TAA-vaccinated mice also received anti-PD-1 and anti-CTLA-4 immune checkpoint blockade (ICB) antibodies twice weekly for two weeks.

**Figure 4.**
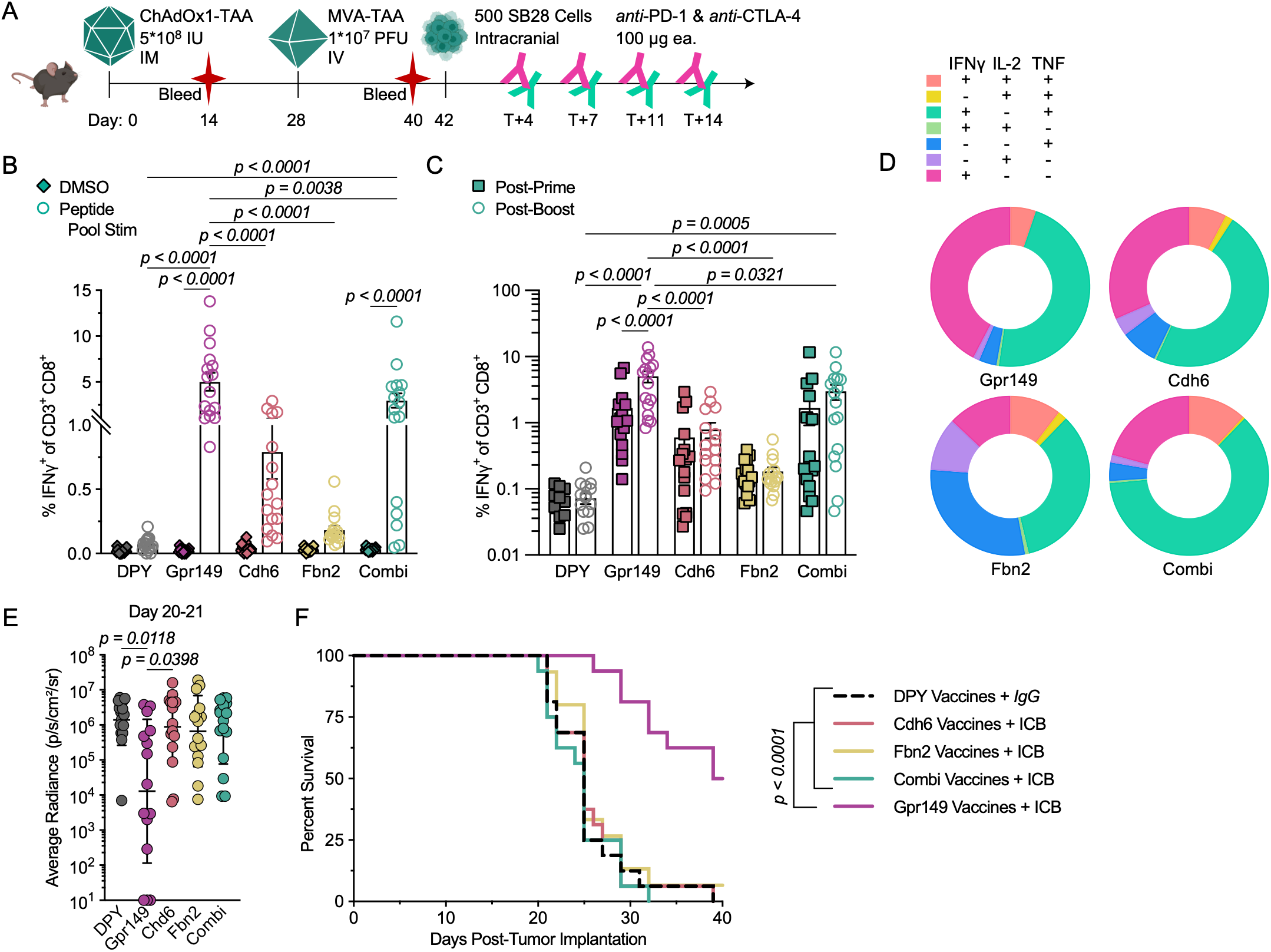
ChAdOx1/MVA-Gpr149 vaccination drives superior immunogenicity and control of intracranial SB28 tumors compared to other TAA vaccines. (**A**) Experimental scheme for (B-F): C57BL/6 mice were vaccinated with ChAdOx1/MVA vaccines encoding SB28 TAAs or DPY one month apart, then challenged intracranially with 500 SB28 tumor cells. Mice received 5x10^8^ IU ChAdOx1 I.M. and 1x10^7^ PFU MVA I.V. All mice receiving SB28 TAA vaccines also received anti-PD-1 and anti-CTLA-4 (ICB; 100 μg each / dose) on days 4, 7, 11, and 14 post-tumor implantation, DPY-vaccinated mice received 200 μg of isotype control. Blood was sampled 14 days post-prime and 12 days post-boost. (**B**) Proportion of CD8^+^ T cells expressing IFN-*γ* after *ex vivo* restimulation of PBMCs with the relevant pool of minimal epitope peptides (*circles*) or without stimulation (DMSO, *diamonds*) 2 weeks post-boost (*n* = 16, pooled from 2 experiments). (**C**) Proportion of CD8^+^ T cells expressing IFN-*γ* after *ex vivo* restimulation of PBMCs with the relevant peptide pool two weeks post-prime (*squares*) or two weeks post-boost (*circles*) (*n* = 16, pooled from 2 experiments). For (B-C), bars depict mean ± SEM, and statistical significance was calculated using 2-way ANOVA with Tukey’s multiple comparisons, with statistically significant comparisons between vaccination conditions shown only between DPY/TAAs and Gpr149/other TAAs for clarity. (**D**) Polyfunctionality of cytokine-expressing CD8^+^ for each vaccination condition post-boost. (**E**) Intracranial IVIS signal for individual mice at D20 or D21. Data are presented as geometric mean ± geometric SD; statistics calculated using a Kruskal-Wallis test with Dunn’s multiple comparisons. (**F**) Survival of prophylactically vaccinated mice; statistics calculated using a log-rank test. All data pooled from 2 independent experiments (*n* = 15-16 mice/group).

We evaluated vaccine immunogenicity in blood post-prime and post-boost via *ex vivo* restimulation of PBMCs with TAA-matched peptide pools. Post-boost, Gpr149- and Combi-vaccinated mice demonstrated significant increases in IFN-*γ*^+^ CD8^+^ T cells after peptide stimulation relative to unstimulated controls (Figure 4B), while responses in Cdh6- and Fbn2-vaccinated mice did not reach statistical significance. Only Gpr149 responses increased significantly from post-prime to post-boost, with a 3-fold mean increase in IFN-*γ*^+^ CD8^+^ T cells (Figure 4C). Prior to tumor implantation, Gpr149 vaccination generated significantly higher frequencies of TAA-specific CD8+ T cells than Cdh6, Fbn2, or Combi vaccination, and only the Gpr149 and Combi vaccines elicited CD8^+^ IFN-*γ* responses significantly greater than DPY-vaccinated mice (Figure 4C). We simultaneously measured TNF and IL-2 induction and found that, post-boost, Combi vaccines induced the highest proportion of polyfunctional CD8^+^ T cells expressing 2 or more cytokines, while Gpr149 vaccines induced the highest proportion of CD8^+^ T cells expressing IFN-*γ* alone (Figure 4D). No TAA-specific responses were detected among CD4^+^ T cells, as was expected given the short peptides used for stimulation (Supplemental Figure 6).

Gpr149-vaccinated mice had a significantly lower tumor burden 3 weeks post-challenge than all other groups as measured by bioluminescence imaging (Figure 4E). Furthermore, the Gpr149-vaccinated mice had significantly prolonged survival, with a median survival of 41 days versus 25 days for all other groups (Figure 4F, Supplemental Figure 7). Survival did not differ between control mice and mice receiving the other TAA vaccines + ICB, indicating that both ICB and low-level TAA-specific immunogenicity were insufficient to protect against SB28. Encouragingly, Gpr149-targeted vaccination drove long-term survival in 50% of mice, demonstrating that peripheral ChAdOx1/MVA vaccination is capable of inducing effective anti-tumor immunity against an immunologically cold intracranial glioblastoma tumor when combined with ICB.

### Gpr149_128-135_ -specific T cells recognize and lyse SB28 tumor cells

Given the strong antigenicity of the Gpr149_128-135_ epitope (VCYNFYTM), we hypothesized that the protective anti-tumor immunity of Gpr149-targeted vaccination *in vivo* was primarily driven by Gpr149_128-135_-restricted recognition of SB28 cells. *In vitro,* we found that Gpr149_128-135_/H-2K^b^-tetramer^+^CD8^+^ (Gpr149_128-135_-specific) T cells but not tetramer^-^CD8^+^ T cells sorted from vaccinated mice expressed IFN*γ* and CD107a when exposed to IFN*γ*-pre-treated wild-type (WT) SB28, but not to untreated or Gpr149-knockout (KO) SB28 (Figure 5A, B). Furthermore, in a real-time growth monitoring assay, Gpr149_128-135_-specific but not tetramer^-^CD8^+^ T cells lysed WT but not Gpr149.KO SB28 in a dose-dependent manner (Figure 5C). Gpr149_128-135_ peptide-pulsing of Gpr149.KO cells restored lysis, demonstrating Gpr149_128-135_-dependent killing of SB28. To assess Gpr149-specific cytotoxicity *in vivo*, Gpr149-vaccinated or naïve mice were infused with splenocytes pulsed with Gpr149_128-135_, Gpr149_447-455_, or no peptide (Figure 5D). Gpr149-vaccinated mice demonstrated ∼100% and ∼25% specific lysis of Gpr149_128-135_- and Gpr149_447-455_-pulsed targets, respectively (Figure 5E, F). Thus, ChAdOx1/MVA-Gpr149 vaccination protects against SB28 tumors primarily through induction of Gpr149_128-135_-specific CD8^+^ T cells mediating antigen-specific cytolysis.

**Figure 5.**
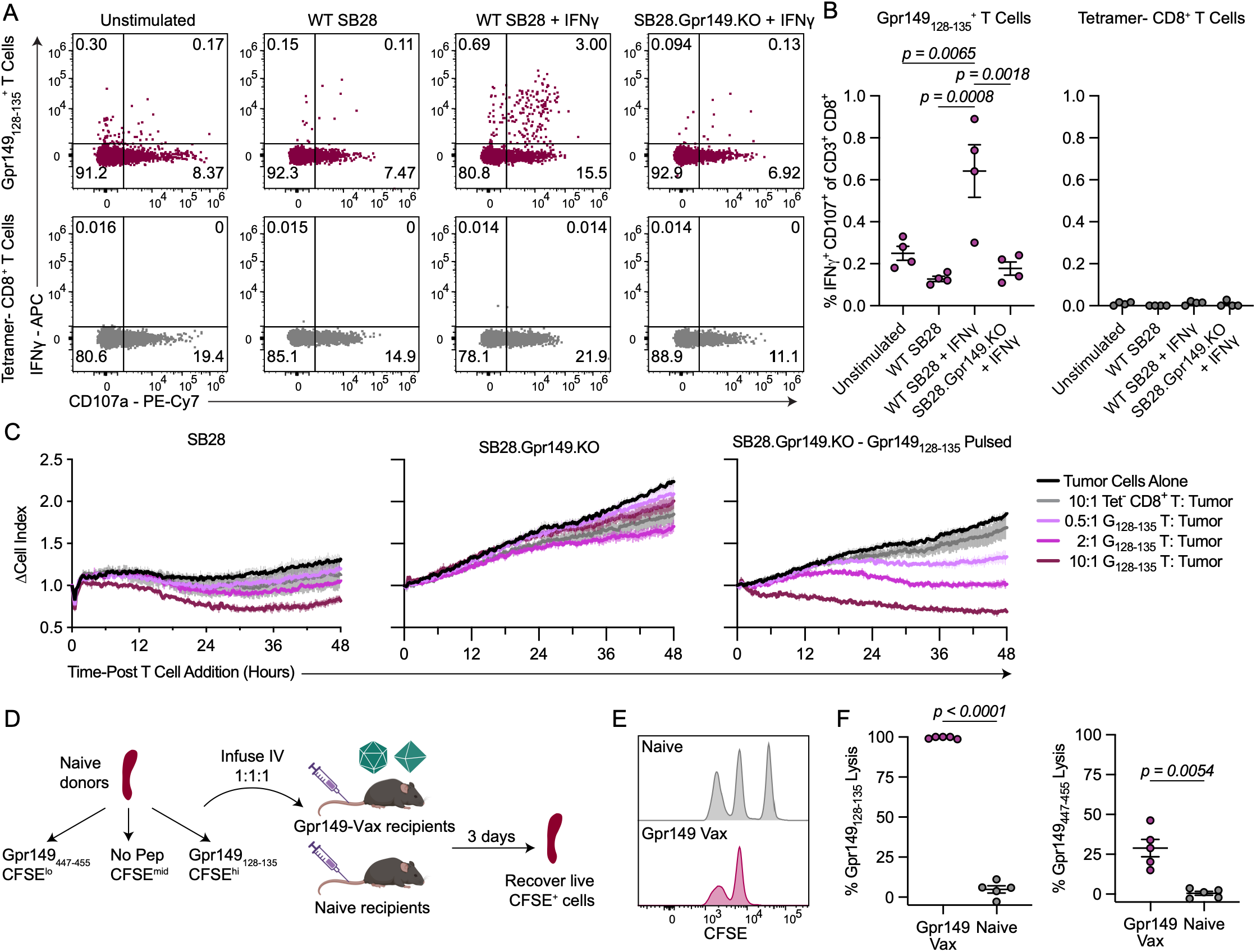
Gpr149_128-135_-specific T cells recognize and lyse SB28 tumor cells. (**A**) Representative reactivity of sorted Gpr149_128-135_-specific or nonspecific T cells against SB28 cells; IFN-*γ* indicates tumor cells pre-treated with IFN-*γ* prior to co-culture. T cells were sorted from the pooled splenocytes of tumor-naïve Gpr149-vaccinated mice six weeks post-boost. (**B**) Proportion of IFN-*γ*^+^CD107a^+^ cells among Gpr149_128-135_-specific or nonspecific T cells after co-culture with SB28 cells (*n =* 4 mice). T cells were sorted independently from splenocytes of individual tumor-naïve Gpr149-vaccinated mice two weeks post-boost. Data are presented as mean ± SEM. Statistical analysis was performed using a 2-way ANOVA with Tukey’s multiple comparisons test; statistics not shown for Gpr149 vs. nonspecific T cell comparisons. (**C**) Fold change over time in cell index (CI) of SB28 cells ascribed in graph titles as measured via a real-time growth monitoring assay during co-culture with sorted Gpr149_128-135_-specific or nonspecific T cells. Thick lines represent mean ∆CI, thin vertical lines represent SEM (*n* = 3 technical replicates). (**D**) Experimental scheme for (E-F): Syngeneic splenocytes from donor mice were pulsed with peptides and labeled with different concentrations of CFSE prior to IV infusion into naïve or Gpr149-vaccinated recipients. (**E**) Representative histograms depicting CFSE signals on splenocytes from Gpr149-vaccinated or unvaccinated mice recovered 3 days after infusion of labeled cells. (**F**) Proportional cell lysis of Gpr149_128-135_ pulsed (*left*) and Gpr149_447-455_ pulsed (*right*) splenocytes by Gpr149-vaccinated (*purple*) or naïve (*grey*) mice (*n* = 5), representative of 2 independent experiments. Data are presented as mean ± SEM; statistical analysis was performed using an unpaired t-test with Welch’s correction.

## Discussion

The present study identifies Gpr149 as an endogenously expressed SB28 antigen and shows that ChAdOx1/MVA vaccination generates Gpr149-specific CD8^+^ T cell responses to mediate protection against intracranial SB28 challenge. Gpr149 was highly expressed in SB28 tumors but minimally detected in immunocompetent adult tissues. Vaccination against Gpr149 elicited an immunodominant CD8^+^ T cell response against Gpr149_128-135_ (VCYNFYTM), with additional responses against Gpr149_447-455_ (RSVFNTITV). Gpr149_128-135_-specific CD8^+^ T cells recognized and lysed wild-type SB28 cells, while recognition was lost after Gpr149 knockout and rescued by exogenous Gpr149_128-135_ peptide loading, demonstrating antigen-dependent recognition of endogenously expressed Gpr149. Together, these findings establish an endogenous antigen system for studying vaccine-induced immunity in the poorly immunogenic SB28 model.

The availability of an endogenous target in the immune checkpoint-refractory SB28 model provides a stringent system for evaluating antigen-targeted immunotherapies. Previous studies of antigen-specific immunity in SB28 have relied largely on transgenic expression of foreign or human antigens. Although these models enable precise tracking of antigen-specific responses, foreign antigens are not subject to the same constraints imposed by self-tolerance and may be expressed at nonphysiological levels. In contrast, endogenous Gpr149 preserves native antigen abundance, processing, and presentation while maintaining tolerance to a self-derived antigen. These features may allow more rigorous assessment of the magnitude and quality of CD8^+^ T-cell responses required for anti-tumor activity in poorly immunogenic tumors. Adoption of this translationally relevant system may aid in rational prioritization of novel therapeutic candidates for clinical testing.

Our antigen discovery strategy further provides a framework for identifying endogenous targets in other preclinical models. Rather than relying exclusively on mutational burden and predicted MHC-I binding, our workflow incorporated tumor-enriched expression, endogenous antigen processing and presentation, immunogenicity verification, and anti-tumor activity. While many pre- and clinical cancer antigen pipelines focus on neoantigen identification, the paucity of neoantigen candidates in the SB28 highlights an important challenge for vaccination against tumors with low mutational burdens. Given the caveats of testing against xenogeneic antigens and/or highly expressed transgenes, identification of endogenous antigens in more tumor models may facilitate improved translatability of preclinical discoveries to human populations.

Alongside other promising new realms of potential cancer antigens such as alternative splice variants^34^ ^35^ and non-canonical peptides derived from the “dark genome”^36–39^, further TAA discovery may lead to identification of more shared cancer antigens that can be targeted with off-the-shelf vaccines, leading to improved outcomes for cancer patients.

In the present study, Gpr149 stood out as the most promising candidate throughout the discovery process. Of all the investigated TAAs, it had the highest ratio of SB28 expression relative to off-target expression and elicited the strongest CD8^+^ T cell responses. Its developmental expression and minimal expression in adult tissues are consistent with a developmentally restricted self-antigen and may have contributed to its favorable combination of tumor specificity and immunogenicity. Meanwhile, the other TAAs showed one or more limiting features, including expression in SB28 only at similar levels to the tested housekeeping gene (*Fbn2, Igf2bp1, Shox2)*, moderate expression in wild-type brain tissue (*Cdh6),* lack of an ability to expand CD8^+^ T cells upon co-culture with TAA-expressing DC2.4 cells (*Fbn2*), and vaccine-induced responses from less than 1% of circulating CD8^+^ T cells on average (*Fbn2, Shox2, Cdh6).* These findings suggest that TAA discovery may be streamlined by applying more stringent thresholds on antigen abundance, endogenous presentation, and immunogenicity throughout the vetting process to quickly narrow down candidates.

The Gpr149 system also revealed potential constraints on antigen targeting. Gpr149 is a G protein-coupled receptor whose ligand is unknown and whose functions remain incompletely characterized, though it has been shown that Gpr149^-/-^ mice have enhanced fertility^40^ and differentiation of oligodendrocyte precursor cells^41^. Indeed, the SB28.Gpr149.KO clone we created had accelerated growth kinetics compared to the paternal line in our real-time growth assay, raising the possibility that immunoediting of Gpr149 could alter tumor cell fitness. While this observation was made using a single knockout clone and would need to be confirmed before attributing this phenotype to Gpr149 loss, antigen-loss variants represent a potential mechanism of immune escape that warrants further investigation in the context of Gpr149-targeted therapy. Furthermore, antigen presentation appeared to represent an important limitation on Gpr149-specific immunity. Gpr149_128-135_ may be processed and presented by SB28 with suboptimal efficiency, as only ∼1-3% of FACS-isolated Gpr149_128-135_-specific T cells were responsive to co-culture with SB28. Moreover, cytokine production and degranulation were observed only after IFN-*γ* pre-treatment of SB28 cells, suggesting that insufficient MHC-I expression may be a critical bottleneck to treating SB28 tumors with antigen-targeted therapies. The low fraction of responsive cells could additionally reflect heterogeneity in T cell functionality or tumor-mediated suppression. While these findings provide a rationale for combining vaccination with agents that enhance tumor MHC-I expression, the clear protective effects of vaccination observed *in vivo* indicate that anti-Gpr149 responses indeed mediate clearance of SB28 tumors.

While we have clearly shown that Gpr149 can be endogenously processed and presented by SB28 cells to facilitate lysis by Gpr149_128-135_-specific CD8^+^ T cells, we acknowledge some limitations to the present study. As the scope of minimal epitope identification was limited to peptides prioritized by the MHC-I binding prediction algorithms, we may have missed additional immunogenic epitopes. However, the immunodominant epitopes eliciting detectable CD8^+^ T cell responses in our assays were among those with the strongest predicted binding, indicating a reasonable correlation between immunogenicity and prediction score. We also did not systematically evaluate CD4^+^ T cell epitopes, and the contribution of Gpr149-specific CD4^+^ T cell responses to the vaccine-mediated anti-tumor immunity remains unclear. Furthermore, the TCR sequences of Gpr149_128-135_-specific CD8^+^ T cell receptors (TCRs) remain unknown. Defining these TCRs could enable future evaluation of Gpr149-targeted adoptive cell therapies. Lastly, while Gpr149 is reported to be upregulated in some human cancers, it does not appear to be upregulated in human glioblastoma. Thus, the principal translation value of Gpr149 in this study is as a preclinical endogenous antigen model rather than as a candidate human glioblastoma target. A number of TAAs, including IL13Rα2, HER2, WT1, gp100, survivin, and MAGE-A1, are reported in glioblastoma and could serve as suitable targets for analogous vaccination strategies^42^.

Our study demonstrates that the ChAdOx1/MVA heterologous prime-boost strategy is well-suited to drive anti-tumor immunity against endogenous glioblastoma antigens. The capacity of these vectors to express long inserts can facilitate CD8^+^ T cell-skewed responses against multiplexed and/or full-length antigens. Moreover, prophylactic vaccination with ChAdOx1/MVA vaccines targeting endogenous Gpr149 effectively reduced intracranial tumor burden and led to long-term survival in a subset of mice. Whether systemic ChAdOx1/MVA vaccination drives durable tumor-specific T cell accumulation within the brain, whether checkpoint blockade is required for efficacy, and whether this approach can therapeutically treat established intracranial tumors remain important questions for future study.

In summary, we demonstrate that an endogenous self-derived antigen in a poorly immunogenic glioblastoma can be targeted by systemic viral-vectored vaccination to generate functional tumor-specific CD8^+^ T cells and protective anti-tumor immunity. Identification of Gpr149 as a novel endogenous SB28 antigen provides a platform to enable further mechanistic evaluation of antigen-targeted therapies in an immunologically stringent model of glioblastoma while avoiding the limitations associated with xenogeneic antigen systems. More broadly, we highlight the potential of systematic identification and validation of endogenous antigens to improve the rigor of preclinical testing. Together, our findings support further evaluation of ChAdOx1/MVA vaccination against glioblastoma, demonstrating a promising new avenue which may lead to therapeutic benefit for cancer patients.

## Methods

### Whole exome sequencing and identification of variants

DNA was extracted from cultured SB28 cells, cultured GL261 cells, and C57BL/6 brain tissue using the Qiagen DNeasy kit according to manufacturer’s protocols. Whole-exome sequencing was performed in duplicate with Illumina MiSeq at the NCI Sequence Core Facility. Bioinformatic analysis was performed using the Illumina DRAGEN pipeline (07.021.595.3.7.5). Variant calling was performed using Dragen Post Somatic Calling Filtering (Illumina) for all pairwise permutations of SB28 vs. brain replicates or GL261 vs. brain replicates. Variants passed if detected in at least two comparisons. The impact of the nucleotide variants was assessed using the Ensembl Variant Effect Predictor (EMBL European Bioinformatics Institute) and passed if the outcome was “High” or “Moderate.” Only those variants with a Variant Allele Frequency of ≥ 30% were considered for neoantigen analysis. Analysis was streamlined using the Mouse nEoanTigen pRedictOr (METRO) pipeline code^31^.

### RNA sequencing and identification of Tumor-Associated Antigen (TAA) candidates

RNA was extracted from cultured SB28 cells, cultured GL261 cells, C57BL/6 brain tissue, and mouse astrocytes using the Qiagen RNeasy kit according to manufacturer’s protocols. RNA-seq was performed in triplicate for each sample by the Center for Cancer Research Sequencing Facility on NovaSeq 6000 SP using Illumina Stranded mRNA Prep and paired-end sequencing. Reads of the samples were trimmed for adapters and low-quality bases using Cutadapt before alignment with the reference genome (mm10) and the annotated transcripts using STAR. Gene expression quantification analysis was performed for all samples using STAR/RSEM tools. Post-alignment QA/QC, quantification to annotation model, and normalization were performed using Partek (Illumina). Only protein-coding and non-mitochondrial genes were included for consideration as TAAs. A student’s t-test comparing the TPM of triplicates for SB28 or GL261 vs. brain or astrocytes was performed to calculate p-values for each gene. The Benjamini-Hochberg procedure was performed on the p-value stacks for each comparison (SB28 vs. brain, GL261 vs. astrocytes, etc.) to calculate q-values and control the false discovery rate. Only those genes with q<0.05 were considered for further analysis. Genes were then further stratified for consideration based on absolute mean TPM (>0.5 in tumor cells, <0.5 in astrocytes and brain).

### RT-qPCR

Tissue samples were collected immediately upon animal euthanasia and snap-frozen in liquid nitrogen. RNA was extracted from samples using the RNeasy kit (Qiagen) according to manufacturer’s instructions. Total cellular RNA (0.5 μg) was used to synthesize first single-strand cDNA using the SuperScript III First-Strand Synthesis kit (ThermoFisher) according to the manufacturer’s protocol. The QuantiTect SYBR Green PCR kit (Qiagen) was used to quantify cDNA by qPCR according to the manufacturer’s instructions. Reactions were run in triplicate on the StepOnePlusTM Real-Time PCR System (Applied Biosystems) at the following conditions: 95°C for 15 mins, (94°C for 15s, 60°C for 30s, 72°C for 30s) x 40 cycles. Gene expression at the mRNA level for each target gene was quantified relative to an internal housekeeping gene control (*Tbp).* To determine relative mRNA expression, mean ΔCt (difference in cycle threshold number) was calculated for each target gene relative to *Tbp* using the formula ΔCt = Ct Target gene - Ct Housekeeping gene. Relative mRNA expression data are shown as 2^-ΔCt^. The primers used are shown in Supplemental Table 7.

### Prediction of MHC binding

For neoepitopes, FASTA sequences corresponding to up to 13 amino acids up- and down-stream of the SNV were input into NetMHC4.1 and the IEDB Consensus Algorithms, which were used to predict 8-14mer epitopes binding to H-2K^b^ and H-2D^b^. For TAAs, FASTA sequences corresponding to the entire amino acid sequence for each gene were input into NetMHC4.1 and the IEDB Consensus Algorithms, which were used to predict 8-11mer epitopes binding to H-2K^b^ and H-2D^b^. All predicted TAA epitopes with an EL Rank < 0.5 (NetMHC4.1) and a Consensus Percentile Rank <1.5 (IEDB) were included for testing in stimulation assays. For prediction of MHC-II binding, Fbn2 minigene sequences were input into NetMHC4.1 and the IEDB Consensus Algorithms, which were used to predict epitopes binding to H-2IA^b^.

### Creation of ChAdOx1/MVA vectors expressing TAAs

The coding sequences for TAA constructs were generated via GeneArt Gene Synthesis (Thermo Fisher Scientific) and codon-optimized for mouse, or were PCR-amplified from SB28 cDNA in the case of Igf2bp1 and Shox2 using the Platinum SuperFi II PCR Master Mix (Invitrogen) and linked together using InFusion cloning (Takara). For all constructs, a sequence coding for the 26 amino acid transmembrane domain of the MHC-II invariant chain was linked to the N terminus of the transgenes. Sequences were inserted into the E1 locus of ChAdOx1 under a CMV immediate early promoter. MVA vectors were constructed with the F11 promoter driving transgene expression. Sequences of cloning vectors were confirmed by Sanger Sequencing (Eurofins Genomics) and alignment in Snapgene. Viruses were produced by the Viral Vector Core Facility in the Pandemic Sciences Institute at the University of Oxford. The purity and identity of the viral vectors were confirmed by PCR.

### Mice

Female 6-8-week-old C57BL/6 and BALB/c mice were purchased from Charles River (USA) or Envigo (UK). Mice received food and water ad libitum and were maintained in 12 hour light/dark cycles. All animal procedures reported in this study that were performed by NCI-CCR affiliated staff were approved by the NCI Animal Care and Use Committee (ACUC) and in accordance with federal regulatory requirements and standards. All components of the intramural NIH ACU program are accredited by AAALAC International. All animal procedures reported in this study that were performed at the University of Oxford were carried out in accordance with the terms of the UK Animals (Scientific Procedures) Act Project License (PPL) (PB050649E).

### Cell culture

The SB28 glioblastoma model was induced via intraventricular transfection of sleeping beauty (SB)-flanked mPDGF, NRasV12, Luciferase, and shRNA silencing of p53; a cell line was established from the tumor^43^. SB28 cells were cultured in DMEM supplemented with 10% fetal bovine serum, 100 U/ml penicillin, 100 μg/ml streptomycin, 2 mM L-glutamine, 100 mM Sodium pyruvate and 50 μM 2-mercaptoethanol. GL261 and HEK293T were maintained in DMEM supplemented with 10% FBS, 100 U/ml penicillin, 100 μg/ml streptomycin and 2mM L-glutamine. DC2.4 cells were maintained in RPMI 1640 supplemented with 10% fetal bovine serum, 100 U/ml penicillin, 100 μg/ml streptomycin, 2 mM L-glutamine, 1% Non-essential amino acids, 0.01 M HEPES Buffer Solution, and 50 μM 2-mercaptoethanol. All adherent cell lines were harvested via trypsination with 0.05% trypsin. T cell media was composed of RPMI 1640 supplemented with 10% FBS, 100 U/ml penicillin, 100 μg/ml streptomycin, 2 mM L-glutamine, 0.01 M HEPES, 100 mM Sodium pyruvate, 1% Non-essential amino acids, 50 μM 2-mercaptoethanol, and 20 U/ml rhIL-2 (Peprotech). All cell lines were maintained at 37°C 5% CO_2_. Cells were negative for mycoplasma contamination. Cell lines were tested negative for pathogen contamination prior to *in vivo* implantation.

### Evaluation of TAA candidate expression in human glioblastoma

RNA-sequencing data (TPM) and clinical annotations for glioblastoma were taken from TCGA-GBM through the Genomic Data Commons. The dataset included 295 tumor samples with RNA-seq data and 617 clinical records. Normal brain cortex expression values were taken from the GTEx dataset (n=270). All tumor samples were confirmed as glioblastoma based on clinical annotations. Nine samples were technical replicates from the same patients. Expression values for duplicates were averaged. After matching expression and clinical data and removing replicates, 286 patients remained for expression analysis. Human orthologs of murine TAAs (SHOX2, FBN2, CDH6, IGF2BP1, and GPR149) were identified using Ensembl gene orthology annotations. For each candidate antigen, fold-change was calculated as the ratio of median glioblastoma expression to median normal brain expression. Prevalence was the percentage of patients with expression above 1 TPM. Tumor expression was compared to the normal tissue median using one-sample t-tests (one-sided, tumor > normal). Multiple testing correction used the Benjamini-Hochberg FDR across five genes, with FDR <0.05 considered significant. Analyses were performed in Python 3.12.10 using pandas v2.3.1, numpy v1.26.4, scipy v1.13.1, and statsmodels v0.14.4. All tests were two-sided unless stated.

### Generation of transgenic DC2.4 cell lines

DC2.4 cells were obtained as a kind gift from Dr. Jeffrey Schlom at NIH. DC2.4 lines expressing each TAA were created via lentiviral transduction. The sequences for Gpr149, Cdh6, Shox2, Igf2bp1, or the 11 Fbn2 minigenes (including spacer sequences) were PCR amplified from the same entry vector plasmids used to create the corresponding ChAdOx1/MVA viral vectors, then inserted into pLVX-TetOne-Puro lentiviral plasmids using InFusion cloning (Takara). Plasmids were verified via Sanger Sequencing before being used for co-transfection of HEK293T-LentiX cells alongside psPAX2 and pMD2.G with X-tremeGENE 9 Transfection Reagent (Sigma). Viral supernatants were collected and used to transduce DC2.4 cells, which were then selected for transgene expression with Puromycin (Gibco). Post-selection, single cells were isolated via FACS to establish monoclonal lines. TAA expression on monoclonal DC2.4 lines was verified via RT-qPCR. Clones expressing each TAA were selected for use in assays. DC2.4 cells were pre-treated for 48 hours with doxycycline (0.1 μg/ml, Sigma) to drive expression of the relevant TAA transgene before use in assays.

### Vaccinations and checkpoint inhibitor treatments

Mice received 5x10^8^ infectious units (IU) of ChAdOx1 virus diluted in PBS injected IM in 50 μl total volume, followed 28 days later by 1x10^7^ plaque forming units (PFU) of MVA virus diluted in PBS injected IV in 100 μl total volume. Checkpoint inhibitors (ICB) were administered intraperitoneally (IP) on days 4, 7, 11, and 14 post-tumor implantation at a dose of 100 μg/dose in a total volume of 100 μl each. Antibodies were purchased from BioXcell (anti-PD-1 clone RPM1-14 or 29F.1A12; anti-CTLA-4 clone 9H10). All non-ICB treated mice received 200 μg of rat IgG2a (Sigma) at ICB administration timepoints.

### Tumor implantations and survival

Intracranial tumors were implanted into mice anesthetized with a combination of xylazine (20 mg/ml) and ketamine (100 mg/ml) diluted in 0.9% NaCl at a 1:1:4 volume ratio, at a total dose of 0.1ml per 20g body weight. 500 SB28 cells in 2 μl of HBSS/DNase (200 μg/ml, Roche) were implanted stereotactically through a 25G Burr hole in the skull using a 33G Hamilton Syringe into the right parietal lobe, 4 mm anterior and 2 mm lateral to the bregma, at a depth of 2 mm, over a period of 10 minutes. Cells were pre-treated in vitro with 100 U/ml of IFN-*γ* (Peprotech) for 48 hours prior to implantation. Mice were euthanized at humane endpoint when body weight fell below 15% of maximum body weight, or when dorsal cranial expansion resulting from tumor outgrowth exceeded a volume of 300 mm^3^ upon measurement with calipers.

### IVIS imaging

*In vivo* tumor measurement was enabled by expression of the luciferase transgene in SB28 cells^16^. D-Luciferin Potassium Salt (GoldBio) was dissolved in PBS (30 mg/ml) prior to use *in vivo*. Mice were injected IP with 100 μl Luciferin 10 minutes prior to imaging with an In Vivo Imaging System (IVIS) Lumina Series III (Revvity). Images were processed using Living Image software (Revvity). Background average radiance was subtracted from the average radiance centered on the head of each mouse. Any negative values resulting from background subtraction were set to 1 for data visualization.

### Processing of blood and splenocytes

Blood was collected retro-orbitally in tubes containing EDTA (12.5 mM, Sigma). Two rounds of red blood cell lysis were performed using a buffer comprised of NH_4_Cl, KHCO_3,_ and tetrasodium EDTA (pH 7.3). Spleens were mashed over a 70 μM strainer prior to 5 minutes of ACK lysis (Quality Biological) and washing with RPMI 1640.

### Surface staining, intracellular cytokine staining, and flow cytometry

Prior to surface staining, cells were incubated for 10 min at 4°C with 5 μg/ml anti-CD16/CD32 (2.4G2, BioLegend) to block Fc receptors, then washed. All tetramers were created by the NIH Tetramer Facility (Emory) and were incubated with cells for 1 hour at 4°C. For surface staining, cells were washed and stained for 20 min at 4°C with viability dye (LIVE/DEAD Aqua, Invitrogen) and fluorescently conjugated mAbs against surface molecules according to different staining panels; anti-CD3-AlexaFlour700 (clone 17A2, BioLegend), anti-CD8-SparkUV387 (clone 53–6.7, BioLegend), anti-CD4-BV421 (clone GK1.5, BioLegend). For intracellular cytokine staining (ICS), cells were stimulated with target cells or peptides (4 mg/ml per peptide unless stated otherwise, Genscript) in the presence of DNAseI (20µg/ml, Roche) and costimulatory anti-CD28 (2 µg/ml, BioLegend) at 37°C for 5 hours, adding brefeldin A (10 µg/ml, BioLegend) for the last 4 hours to allow accumulation of intracellular cytokines. Cells were then surface stained, fixed and permeabilized (BD Cytofix/Cytoperm), then stained intracellularly for cytokine production with anti-IFN-*γ*-APC (clone XMG1.2, BioLegend), anti-IL-2-PE (clone JES6-5H4, BioLegend), and anti-TNF-FITC (clone MP6-XT22, BioLegend), then acquired on an Aurora spectral flow cytometer (Cytek). Data were analyzed with FlowJo software v10 (Tree Star, Ashland, Oregon). Analysis of multifunctional CD8^+^ T-cell responses was performed via a Boolean analysis of IFN-*γ*^+^, TNF^+^, and IL-2^+^ events in the CD8^+^ gate using FlowJo. Pestle (NIH, Bethesda) and SPICE (Vaccine Research Centre, NIH, Bethesda) software were used to generate graphical representations of proportions of T cells expressing 1, 2, or all 3 cytokines^44^.

### RMA-S MHC-I binding assay

RMA-S cells were obtained as a generous gift from Dr. David Margulies at NIH. RMA-S cells were cultured in the presence of human β2-microglobulin (5 μg/ml, Sigma) and increasing concentrations of peptides (85% purity, Genscript) for 16 hours. Cells were then stained simultaneously with fluorescently conjugated H-2D^b^ (KH95, BioLegend) and H-2K^b^ (AF6-88.5, BioLegend) antibodies prior to flow cytometry.

### Generation of SB28.Gpr149.KO cell line

TrueGuide Synthetic gRNA targeting Gpr149 (AAGUCAUAAUUCUACGGAGA, ThermoFisher), TrueCut Cas9 Protein v2 (ThermoFisher), and Lipofectamine CRISPRMAX (ThermoFisher) were combined in Opti-MEM (Gibco), then added to SB28 cells for 24 hours. Single cells were plated into 96-well plates by flow-assisted cell sorting (BD FACS Aria) to establish clones. To screen for complete Gpr149 knockout, DNA was isolated from each clone, the genetic region flanking the cut site was amplified via PCR, and the PCR product was purified via agarose gel electrophoresis prior to Sanger Sequencing. Knockout efficiency was measured with the SeqScreener Gene Edit Confirmation App (ThermoFisher). One monoclonal line with complete Gpr149 knockout was selected for use in all experiments.

### DC2.4 co-culture assay

Mice were primed IM with 5x10^8^ IU with a given ChAdOx1-TAA vaccine and boosted 28 days later with 1x10^8^ IU of the same ChAdOx1-TAA vaccine IM. Splenocytes were isolated 2 weeks post-boost. To expand TAA-specific CD8^+^ T cells, splenocytes pooled from 3-5 vaccinated mice were co-cultured with irradiated DC2.4 lines expressing the relevant TAA transgene in the presence of human recombinant IL-2 (20 U/ml, Roche). Splenocytes from mice vaccinated with ChAdOx1-Combi were separately stimulated with either DC2.4.Shox2 or DC2.4.Igf2bp1. After 9 days, T cells were restimulated with the same cognate irradiated DC2.4-TAA cells or WT DC2.4 cells for 12 hours in the presence of anti-CD28 (2 µg/ml, BioLegend) and DNase (20 µg/ml, Roche), then for an additional 10 hours in the presence of Brefeldin A (10 µg/ml, BioLegend). IFN-*γ* secretion was measured by ICS and flow cytometry.

### Stimulation assay of tetramer-labeled T cells

T cells were sorted from the spleens of ChAdOx1/MVA-Gpr149 vaccinated mice via FACS after staining with Gpr149_128-135_/H-2K^b^ and Gpr149_447-455_/H-2D^b^ tetramers and surface staining antibodies. Control CD3^+^CD8^+^ T cells were negative for both tetramers. For the data shown in Figure 5A, T cells were FACS-isolated from the pooled splenocytes of tumor-naïve mice (*n* = 3) vaccinated according to the prophylactic ChAdOx1/MVA-Gpr149 scheme six weeks post-boost. For the data shown in Figure 5B, T cells were sorted individually from splenocytes of individual tumor-naïve mice vaccinated according to the prophylactic ChAdOx1/MVA-Gpr149 scheme two weeks post-boost. T cells were rested overnight at 37°C in T cell media supplemented with 20 U/ml IL-2 prior to use in the assay. Some target SB28 cells or SB28.Gpr149.KO cells were treated with rm-IFN*γ* (100 U/ml, Peprotech) for 48 hours prior to use in assay. 25,000 T cells were co-cultured with 50,000 tumor cells for 5 hours in T cell media in round-bottom 96-well plates in the presence of IL-2 (20U/ml), DNase (20 μg/ml, Roche), anti-CD28 (2 μg/ml, BioLegend), and CD107a-PE-Cy7 (1D4B, 125 ng/ml, BD). Brefeldin A (5 μg/ml, BioLegend) was added for the last four hours. ICS and flow cytometry was then performed as described above.

### Real-time monitoring of cell growth

Target SB28 cells or SB28.Gpr149.KO cells were treated with rm-IFN-*γ* (100 U/ml, Peprotech) for 48 hours prior to harvest. Upon harvesting, cells for some test conditions were pulsed with Gpr149 peptides (4 μg/ml, 85% purity, Genscript) for 2 hours at 37°C, then washed thoroughly with PBS. Non-pulsed cells were handled identically. 2500 cells were plated in 100 μl T cell media supplemented with IL-2 (20 U/ml) in 16-well E-Plates (Agilent) and electrical impedance was measured every 15 minutes using an XCELLigence Real-Time Cell Analysis MP instrument (Agilent). A total of 25,000, 5000, or 1250 T cells in a volume of 100 μl T cell media were added 24 hours after initial plating of tumor cells. T cells were derived from the splenocytes of tumor-naïve mice vaccinated according to the prophylactic ChAdOx1/MVA-Gpr149 regimen two weeks post-boost. T cells were FACS-isolated after staining with anti-CD3 (17A2, BioLegend), anti-CD8 (53-6.7, BioLegend), and Gpr149_128-135_/H-2K^b^- and Gpr149_447-455_/H-2D^b^-tetramers. Tetramer-negative control T cells were negative for both tetramers. T cells were rested overnight at 37°C in T cell media supplemented with 20 U/ml IL-2 prior to use in the assay. ∆Cell Index was calculated by the XCELLigence Real-Time Cell Analysis software (Agilent) from the time point directly prior to the addition of T cells.

### In vivo lysis assay

Splenocytes from naïve C57BL/6 mice were harvested and split three ways prior to pulsing for 2 hours with Gpr149_128-135_ or Gpr149_447-455_ peptides (10 μg/ml, Genscript). Cells were then labeled with 1 μM, 0.33 μM, or 0.11 μM CFSE (ThermoFisher) (CFSE^hi^: Gpr149_128-135_ pulsed, CFSE^mid^: non-pulsed, CFSE^lo^: Gpr149_447-455_ pulsed, respectively), mixed at a 1:1:1 ratio in PBS, and 2x10^6^ cells/condition were infused IV into vaccinated or naïve recipients. Vaccinated recipients were treated with ChAdOx1/MVA-Gpr149 according to the prophylactic dosing scheme prior to the infusion of CFSE-labeled cells 6 weeks post-boost. 72 hours post-infusion, spleens were harvested from recipients and run on a flow cytometer without additional staining. Percent lysis was calculated as (#Non-pulsed - #Pep-Pulsed)/#Non-Pulsed x 100.

### Statistical analysis and figure preparation

Statistical analysis was performed using GraphPad PRISM (V10) or Python. Figures were prepared using Adobe Illustrator, FlowJo, SPICE, GraphPad PRISM, and Biorender.

## Acknowledgments

We thank Ferenc Livac, Shafiuddin Siddiqui, and Kathy McKinnon at the Center for Cancer Research Flow Cytometry Core (NIH); Claire Powers at the Pandemic Sciences Institute Viral Vector Core Facility (University of Oxford); V. Clark and H. Gray at the Functional Genomics Facility (University of Oxford), and Devorah Gallardo at the Building 37 Animal Facility (NIH) for technical assistance. We thank the NIH Tetramer Core Facility (contract number 75N93020D00005) for providing all tetramers used in this manuscript. CSKL thanks the Swiss National Science Foundation and Academy of Medical Sciences Springboard Award for support.

## Funding

This research was supported in part by the Center for Cancer Research, National Cancer Institute, National Institutes of Health Intramural Research Program project number ZIA BC 011877 and federal funds from the National Cancer Institute, National Institutes of Health, under contract HHSN261200800001E. The contributions of the NIH authors are considered works of the United States Government. The findings and conclusions presented in this paper are those of the authors and do not necessarily reflect the views of the NIH or the U.S. Department of Health and Human Services.

This work was supported in part by the Ludwig Institute for Cancer Research. EES was supported by a Marshall Scholarship from the Marshall Aid Commemoration Commission and the NIH-Oxford-Cambridge-Scholars Program. HO was supported by NIH grant 1R35NS142982.

## Author contributions

EES, CSKL, BJVdE, and MT conceived and designed the experiments. EES, TH, LL, SAM, JM, VPA, NB, MZ, WZ, DD, HS, and HK performed the experiments. EES, TH, and MB analyzed data. BJVdE and MT acquired project funding. HO and MG contributed resources. CSKL, BJVdE, and MT supervised the study. EES administered the project, performed data visualization, and wrote the original manuscript draft. All authors reviewed and edited the manuscript.

## Competing interests

BJVdE and CSKL are inventors on a patent that covers viral vectors and methods for the prevention and treatment of cancer.

## Data and materials availability

All data associated with this study are available in the main text or the Supplementary Materials. Processed RNA-seq and WES data are available in an online database: https://zenodo.org/records/21416624. ChAdOx1 and MVA vaccines can be provided via a material transfer agreement to academic researchers submitting a research proposal to University of Oxford corresponding authors CSKL or BVDE. SB28 and SB28-Gpr149.KO cells can be provided via a material transfer agreement to academic researchers submitting a research proposal to National Institutes of Health corresponding author MT. Requests must be evaluated for biosafety and regulatory compliance, so any transfer of materials is subject to review and approval by the University of Oxford or the National Institutes of Health. Transfer of materials requires the execution of appropriate legal agreements. All other materials used or generated in this study are commercially available or will be supplied upon reasonable request.

---

**Figure S1.**
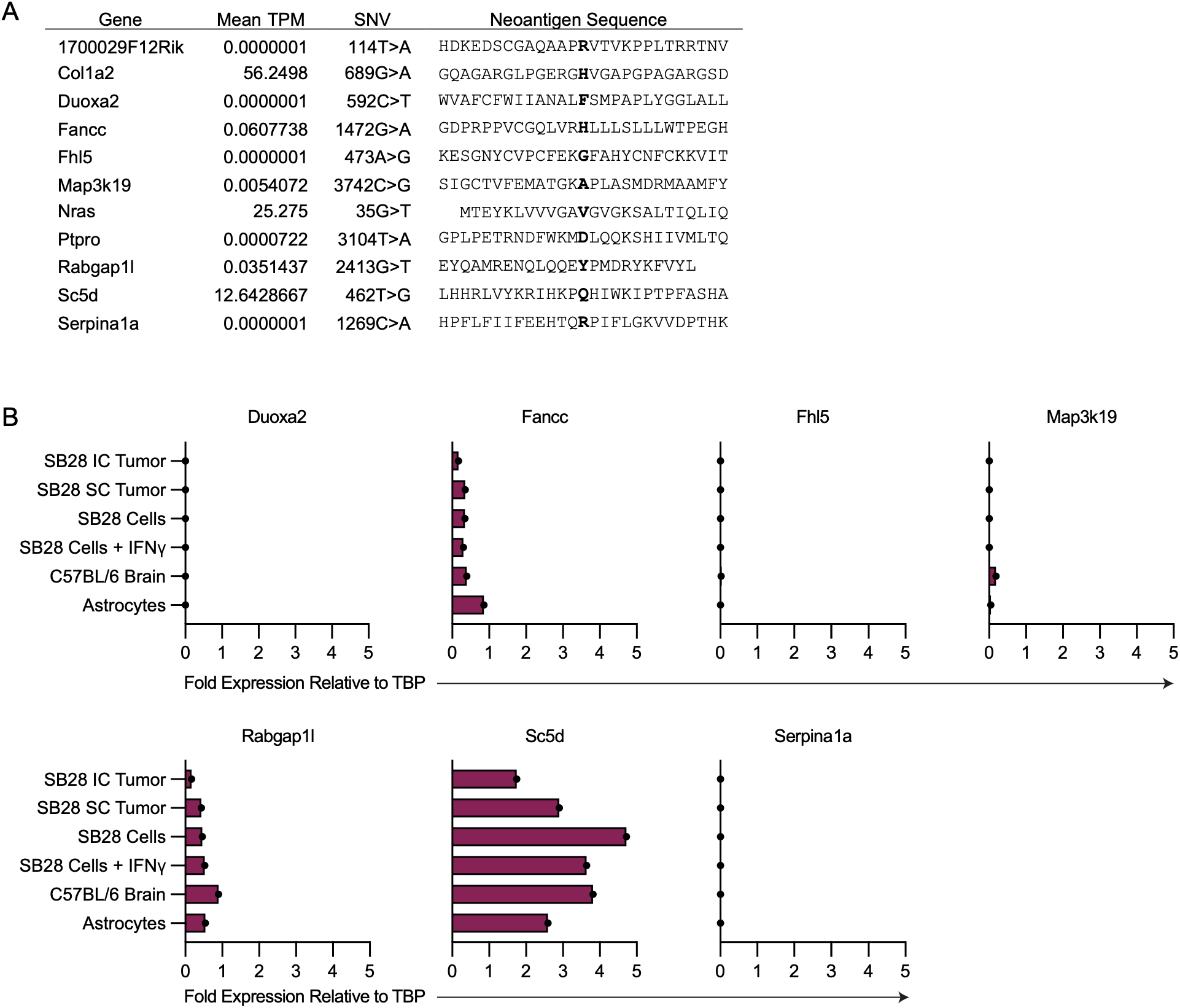
Expression levels of SB28 Neoantigen candidates. (**A**) Expression of all SB28 neoantigen candidates based on RNA-seq of SB28 cell line. Values are mean expression of triplicates. TPM, transcripts per kilobase million. (**B**) Fold change in neoantigen candidate gene expression in various SB28 samples or control tissues relative to housekeeping gene as measured by RT-qPCR. Dots represent unique tissue samples from individual mice (*n =* 1/test condition). TBP, TATA-Box Binding Protein; IC, intracranial; SC, subcutaneous.

**Figure S2.**
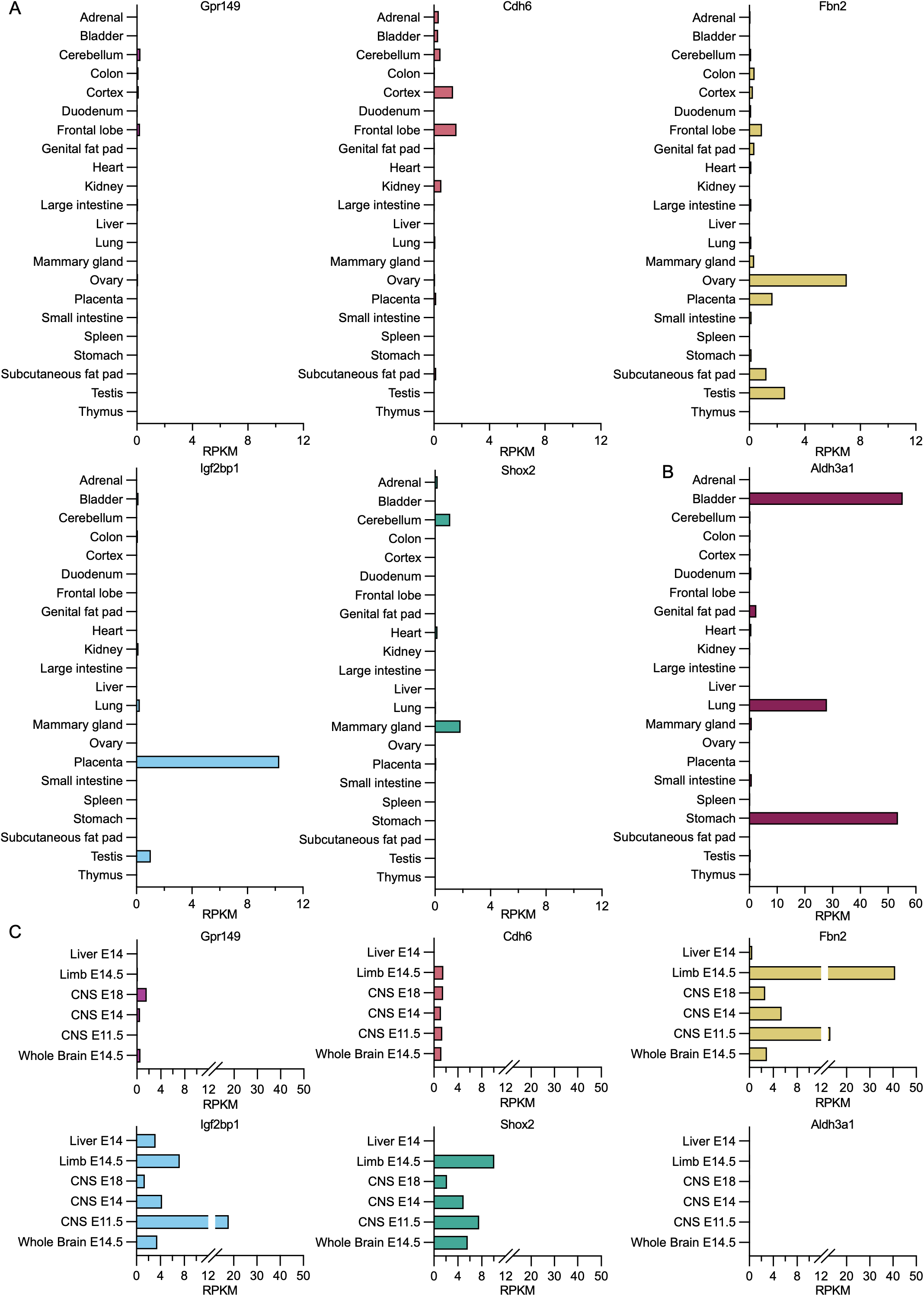
Top TAA candidates are minimally expressed in adult immunocompetent tissues. (**A**) Transcriptomic expression of top 5 SB28 TAA candidate genes in various murine adult tissues in the Mouse ENCODE transcriptome dataset from NCBI Gene. RPKM, reads per kilobase million. (**B**) Transcriptomic expression of the example TAA candidate gene disqualified from further consideration based on expression in various murine adult tissues based on the mouse ENCODE transcriptome dataset from NCBI Gene. (**C**) Transcriptomic expression of selected genes in various murine embryonic tissues from the Mouse ENCODE transcriptome dataset from NCBI Gene. Number following “E” designates age of embryo in days.

**Figure S3.**
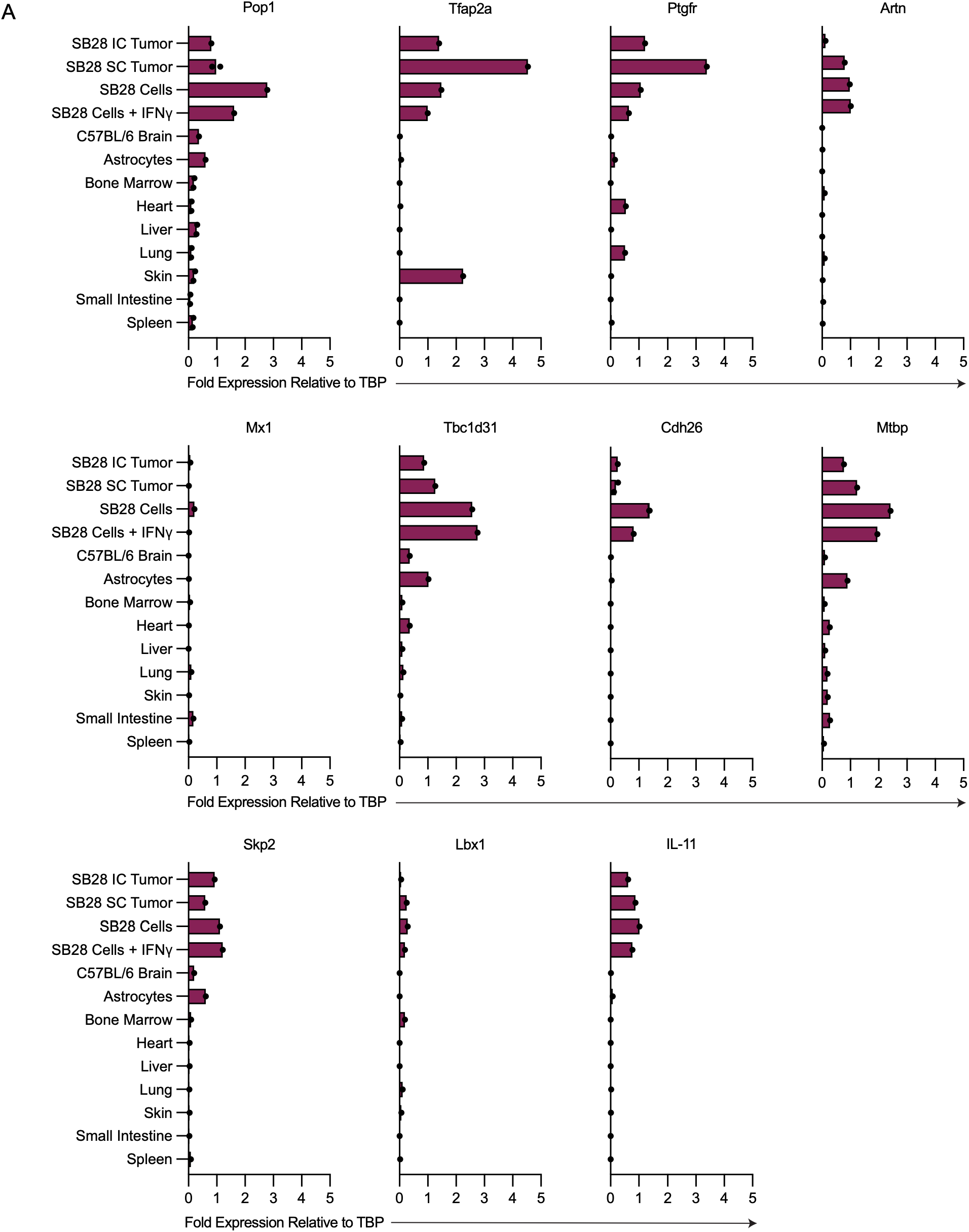
TAA candidates excluded from further consideration based on low measured SB28 expression and/or high expression in peripheral tissues. (**A**) Fold change in TAA candidate gene expression in various SB28 samples or peripheral tissues relative to the housekeeping gene as measured by RT-qPCR. Dots represent unique tissue samples from individual mice (*n =* 1-2/test condition). TBP, TATA-Box Binding Protein; IC, intracranial; SC, subcutaneous.

**Figure S4.**
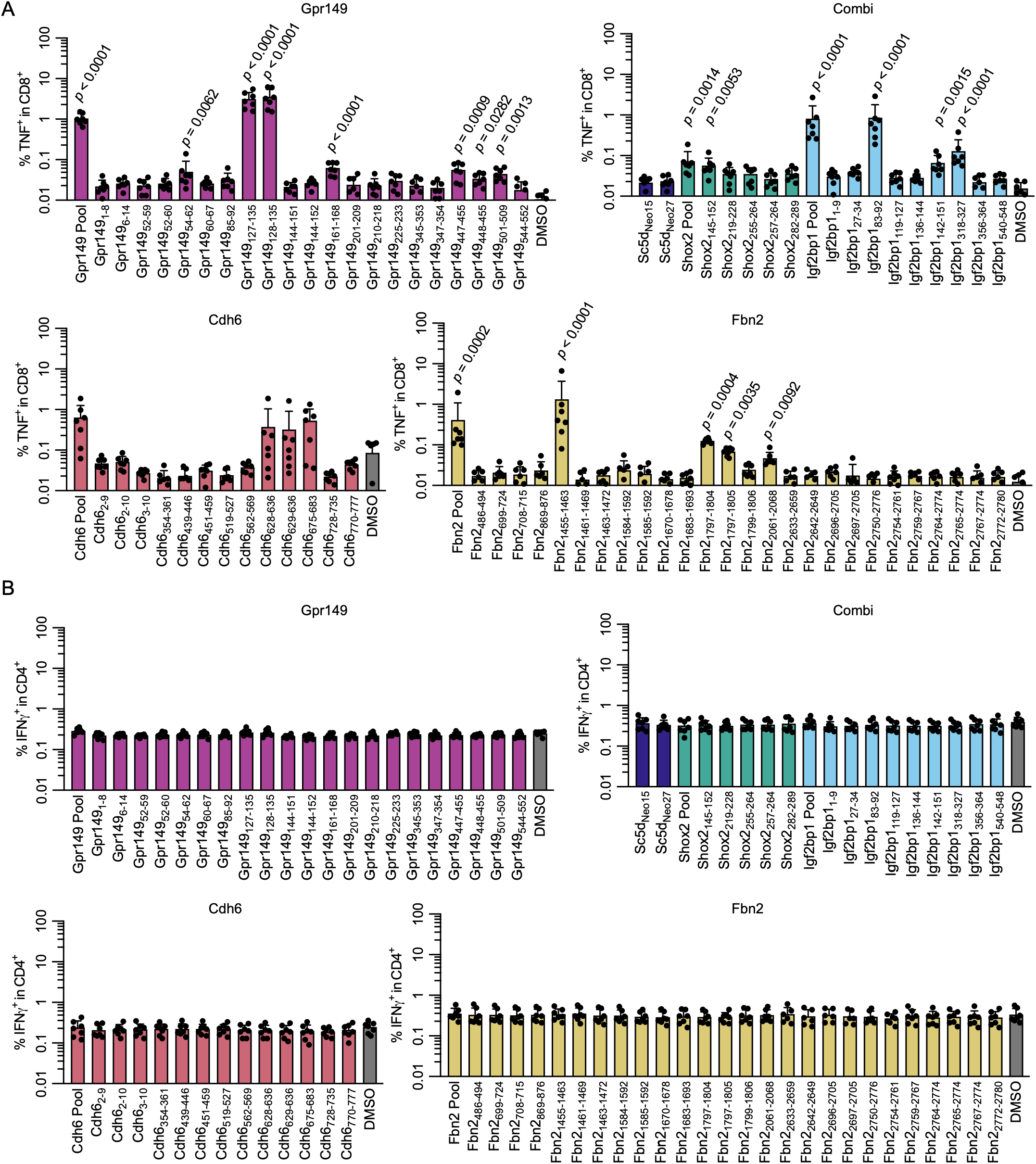
Minimal immunogenic epitopes stimulate TNF expression by CD8^+^ T cells but do not stimulate CD4^+^ cells. (**A**) Frequency of CD8^+^ T cells expressing TNF or (**B**) CD4^+^ T cells expressing IFN-*γ* after *ex vivo* restimulation of splenocytes from vaccinated mice with minimal epitope peptides (4μg/ml). Peptide pools include all peptides for a given gene (4μg/ml each). Dots represent individual mice (*n =* 7/group); pooled from 2 independent experiments. Statistical significance was calculated using a Kruskal-Wallis test with Dunn’s multiple comparisons for each stimulation condition vs. DMSO controls.

**Figure S5.**
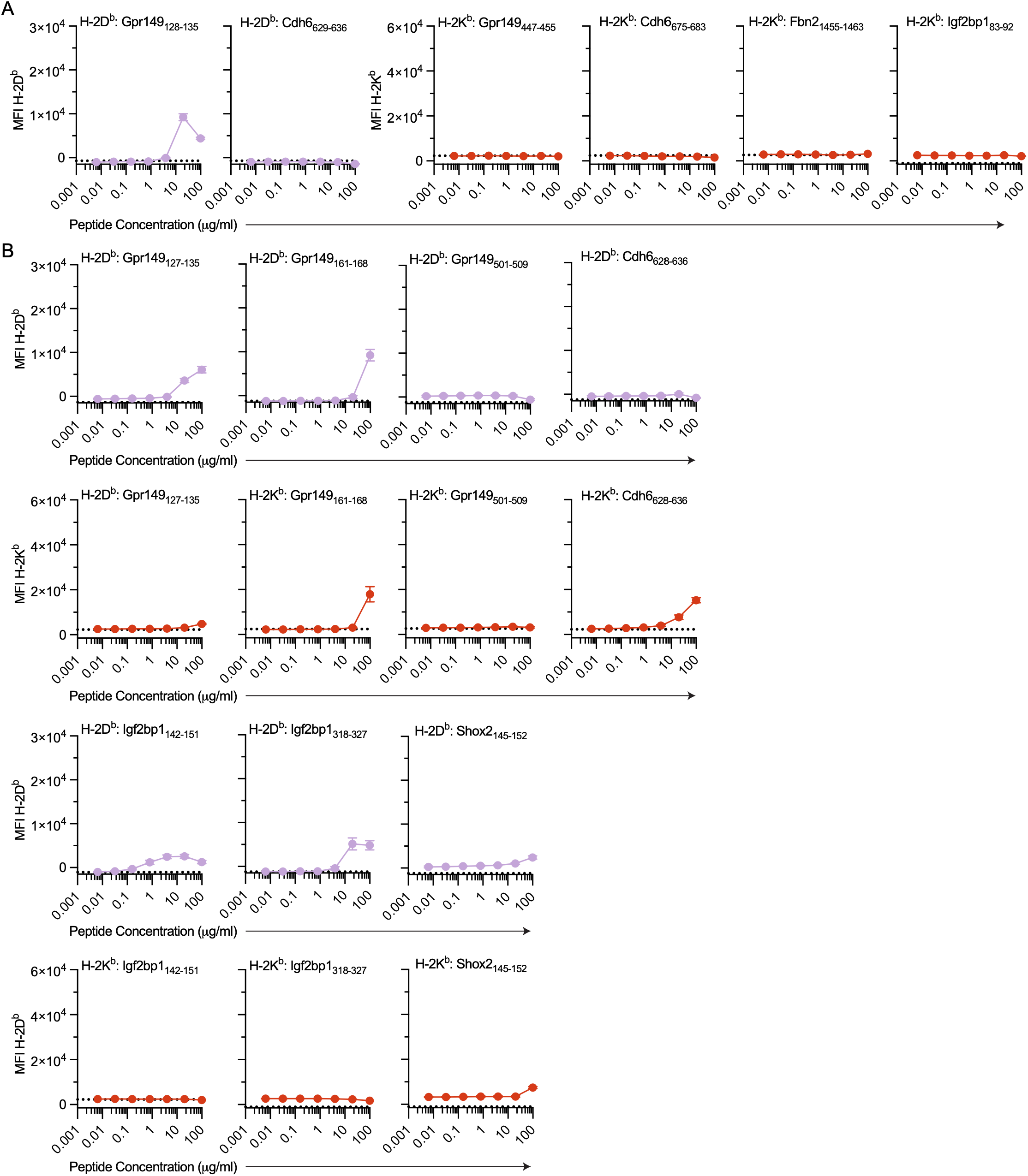
Minimal TAA epitopes mediating insufficient stabilization of H-2K^b^ or H-2D^b^ on RMA-S cells. Median fluorescence intensity (MFI) of surface H-2D^b^ or H-2K^b^ on RMA-S cells measured by flow cytometry after overnight culture in the presence of peptide at increasing concentrations. (**A**) Plots showing expression of H-2D^b^ or H-2K^b^ for non-predicted MHC-I allele for epitopes shown in Figure 3B. (**B**) Plots showing expression of H-2D^b^ or H-2K^b^ for all other tested minimal epitopes. Error bars represent standard deviation of triplicates; representative of 2 independent experiments.

**Figure S6.**
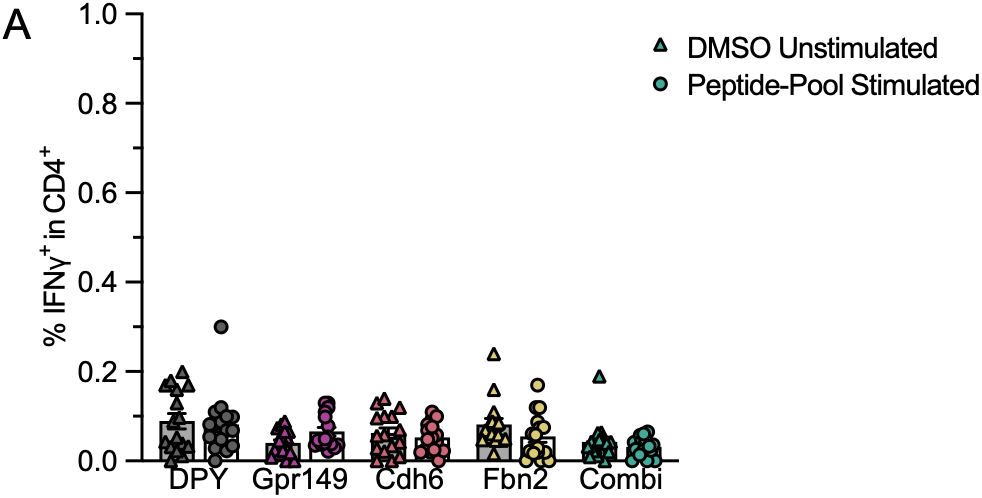
Restimulation of PBMCs from vaccinated mice with relevant TAA peptide pools did not elicit IFN-*γ* expression by CD4^+^ T cells. Mice were vaccinated with ChAdOx1/MVA-TAA or DPY vaccines and blood was sampled 2 weeks post-boost. (**A**) Proportion of CD4^+^ T cells expressing IFN-*γ* after *ex vivo* restimulation of PBMCs with the relevant peptide pool. PBMCs from combi-vaccinated mice were stimulated with pooled Igf2bp1, Shox2, and Smc2 peptides; PBMCs from DPY (irrelevant antigen)-vaccinated mice were stimulated with all experimental peptides (i.e., Gpr149, Cdh6, Fbn2, Igf2bp1, Shox2, and Smc2). Points represent IFN-*γ* expression for CD8^+^ T cells from individual mice (Triangles, DMSO unstimulated controls; Circles, peptide-pool stimulation). Bars depict mean ± SEM; statistical significance was calculated using 2-way ANOVA with Sídǎk’s multiple comparisons test (all *n.s.*); pooled from 2 independent experiments.

**Figure S7.**
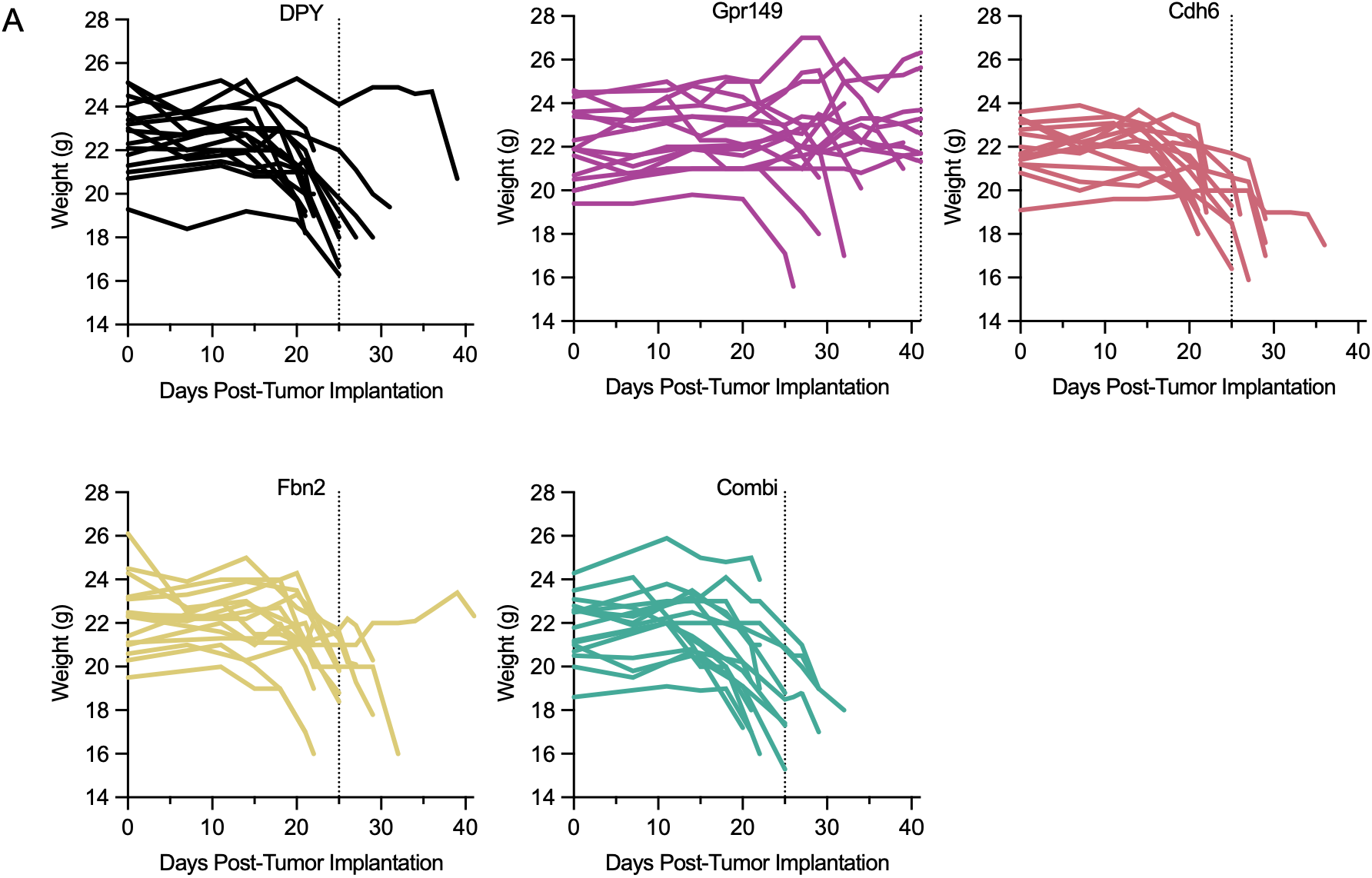
Body weights representing decline in physical condition of mice bearing intracranial SB28 tumors. (**A**) Line graphs depicting change in body weight of mice implanted intracranially with SB28 tumors for each vaccination condition for the experiment described in Fig. 4. Vertical dotted lines represent median survival for each group.

**Supplemental Table 1.**
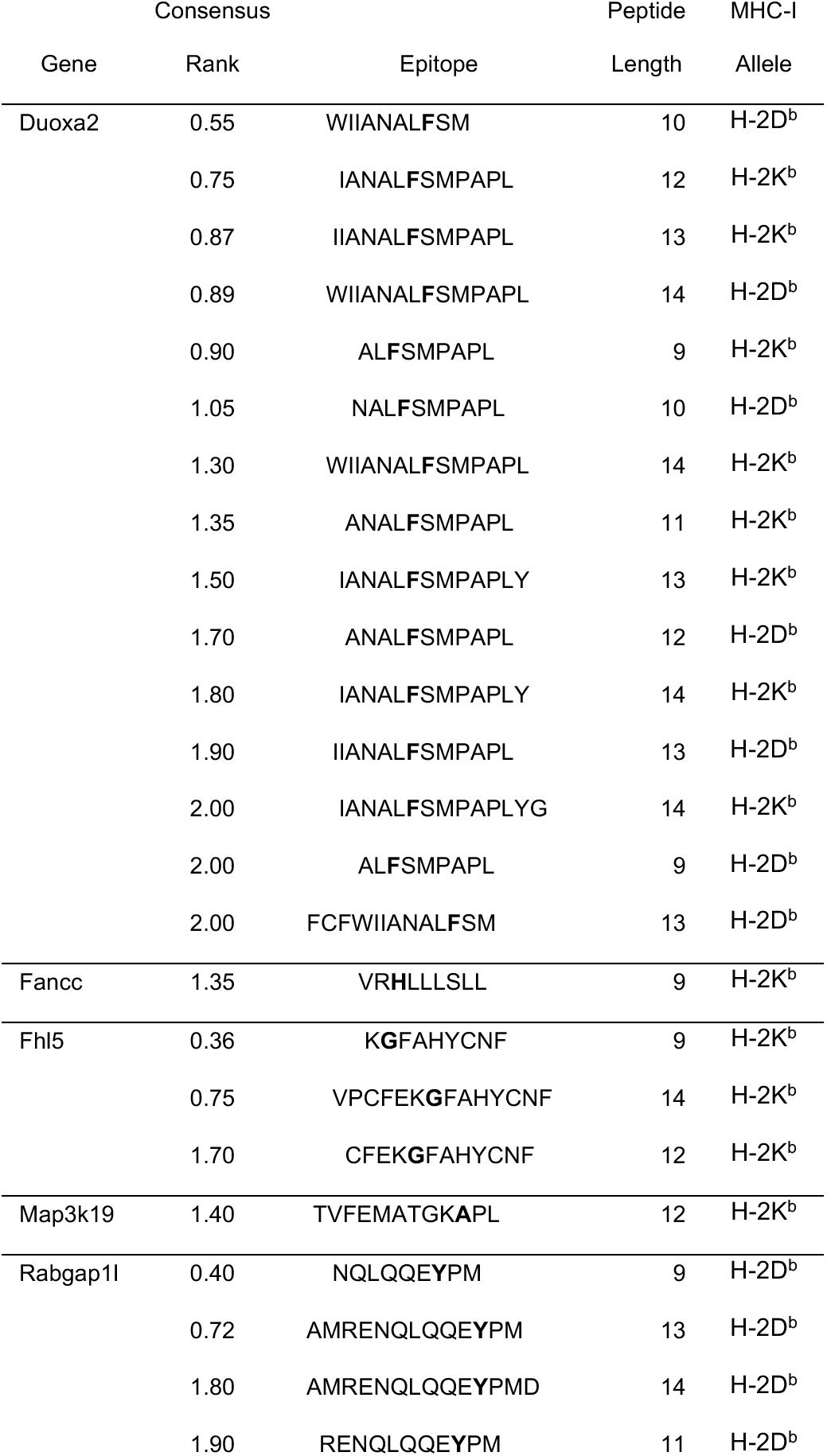
IEDB prediction scores for SB28 neoantigen candidates. 8-14mer minimal epitopes predicted by the IEDB Consensus algorithm to bind H-2D^b^ or H-2K^b^ with Consensus Percentile Rank ≤ 2.0.

**Supplemental Table 2.**
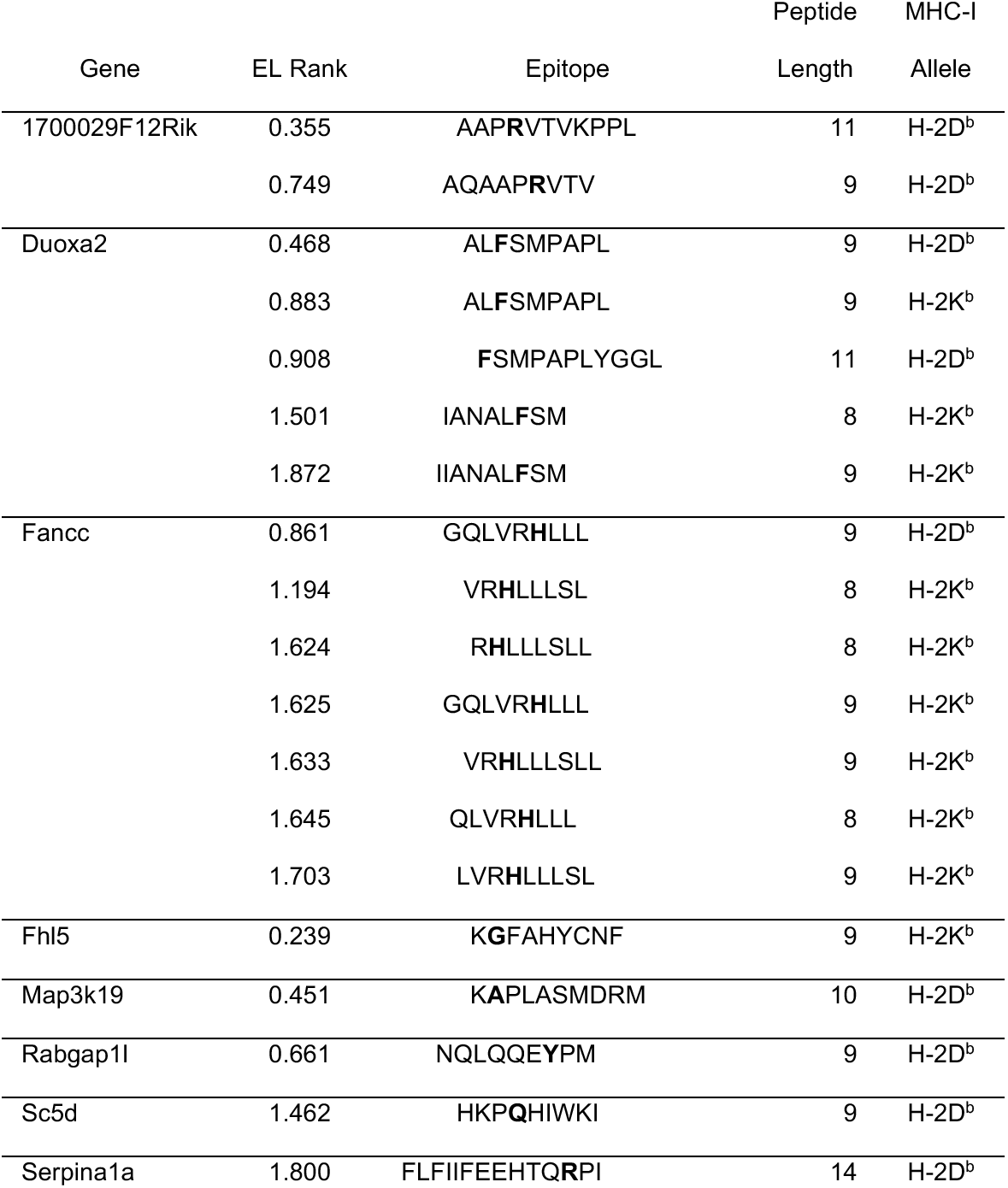
NetMHC4.1 prediction scores for SB28 neoantigen candidates. 8-14mer minimal epitopes predicted by the NetMHC4.1 algorithm to bind H-2D^b^ or H-2K^b^ with EL Rank ≤ 2.0.

**Supplemental Table 3.**
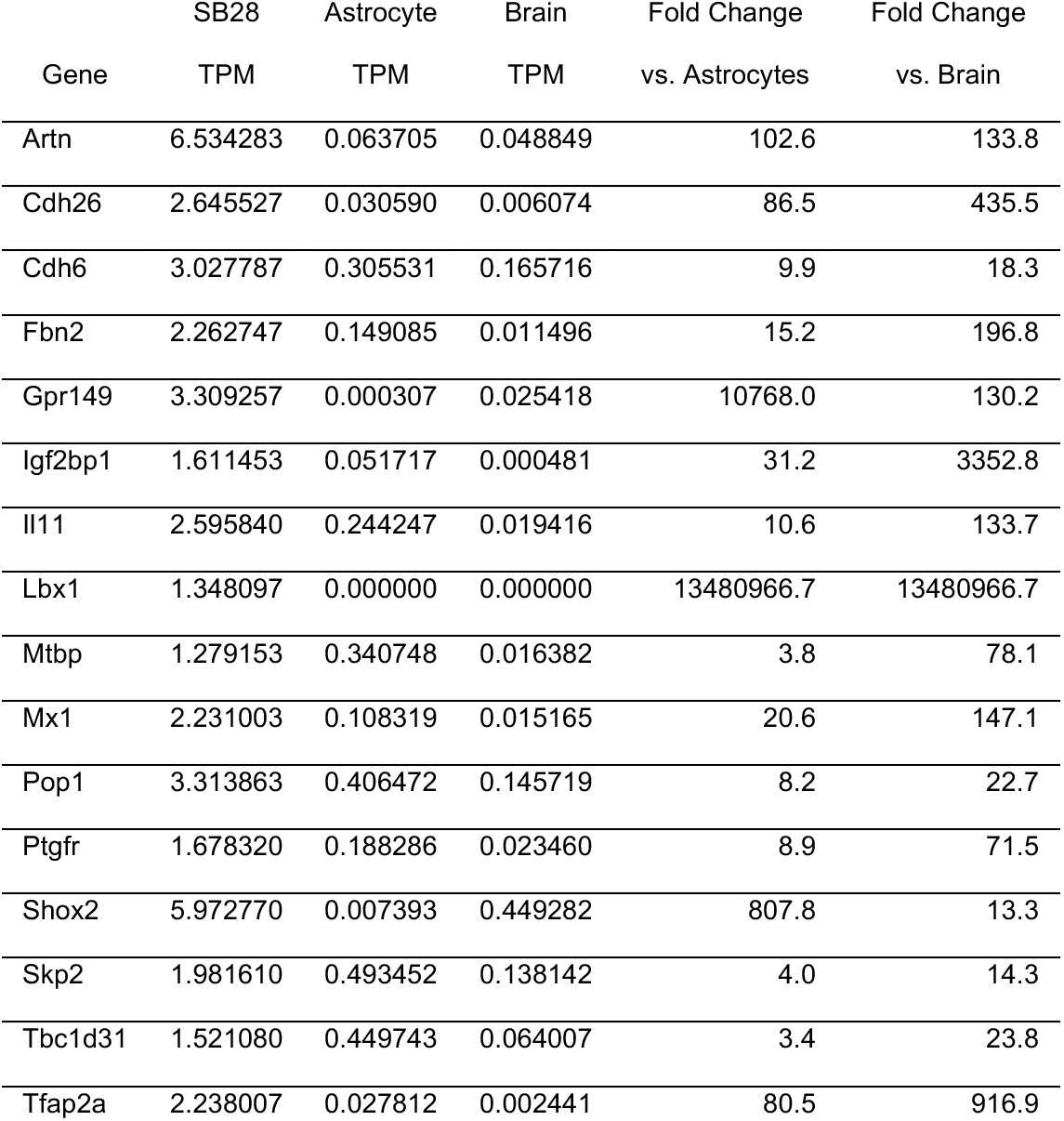
Expression of top 16 SB28 TAA candidates by RNA-seq. RNA from SB28 cells, astrocytes, or C57BL/6 brain was sequenced in triplicate. Expression level for each gene is shown by mean TPM (transcripts per million). Fold change was calculated as mean SB28 TPM / mean astrocyte or brain TPM.

**Supplemental Table 4.**
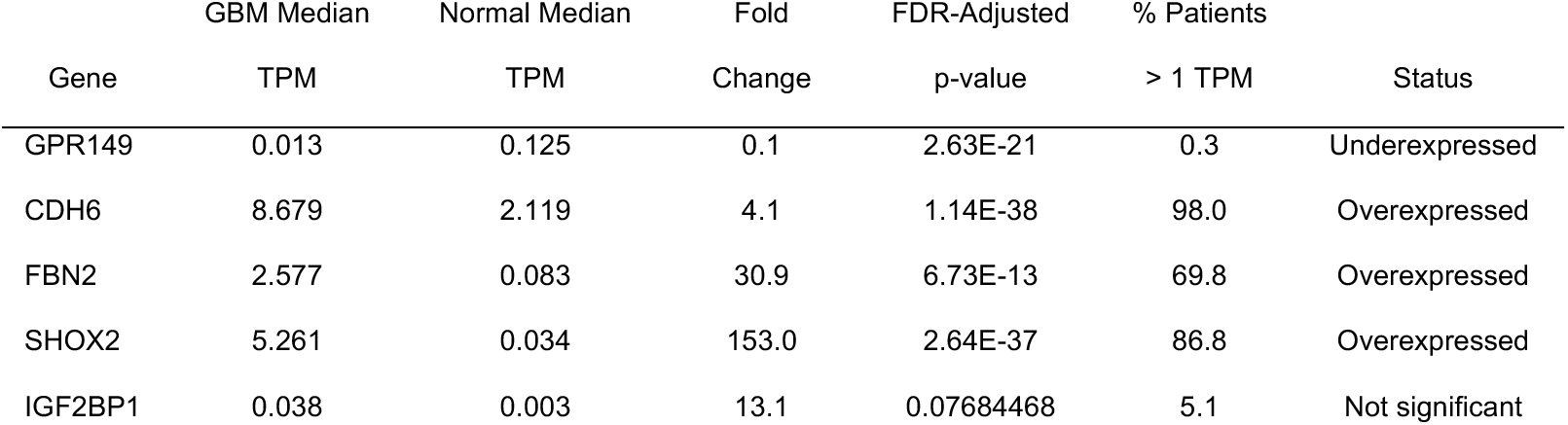
Expression of top 5 SB28 TAA candidates in human glioblastoma. Expression of candidate genes in human IDH mutant glioblastoma (*n =* 295) compared to healthy cortical tissue (*n* = 270), based on TCGA and GTEx datasets, respectively.

**Supplemental Table 5.**
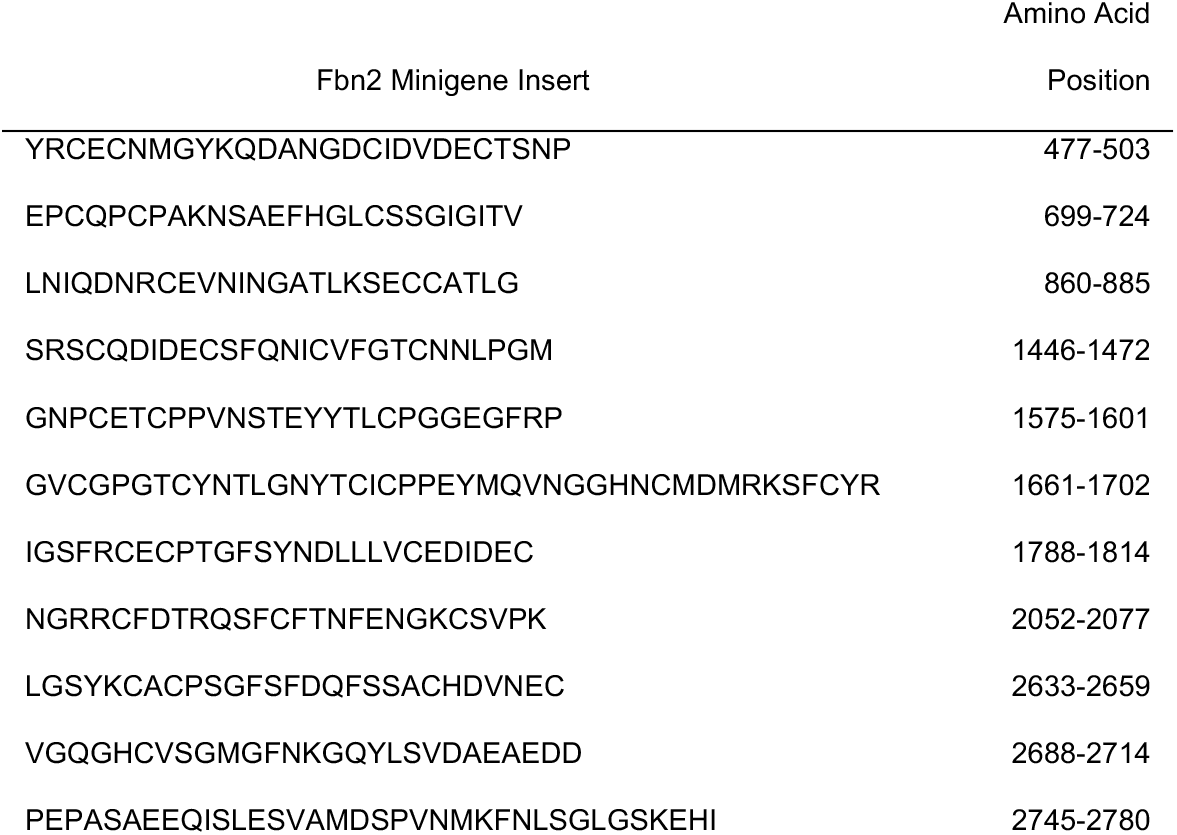
Minigene inserts included in the Fbn2 vaccine constructs. Amino acid sequence of Fbn2 minigenes and position of minigene within whole Fbn2 gene is shown.

**Supplemental Table 6.**
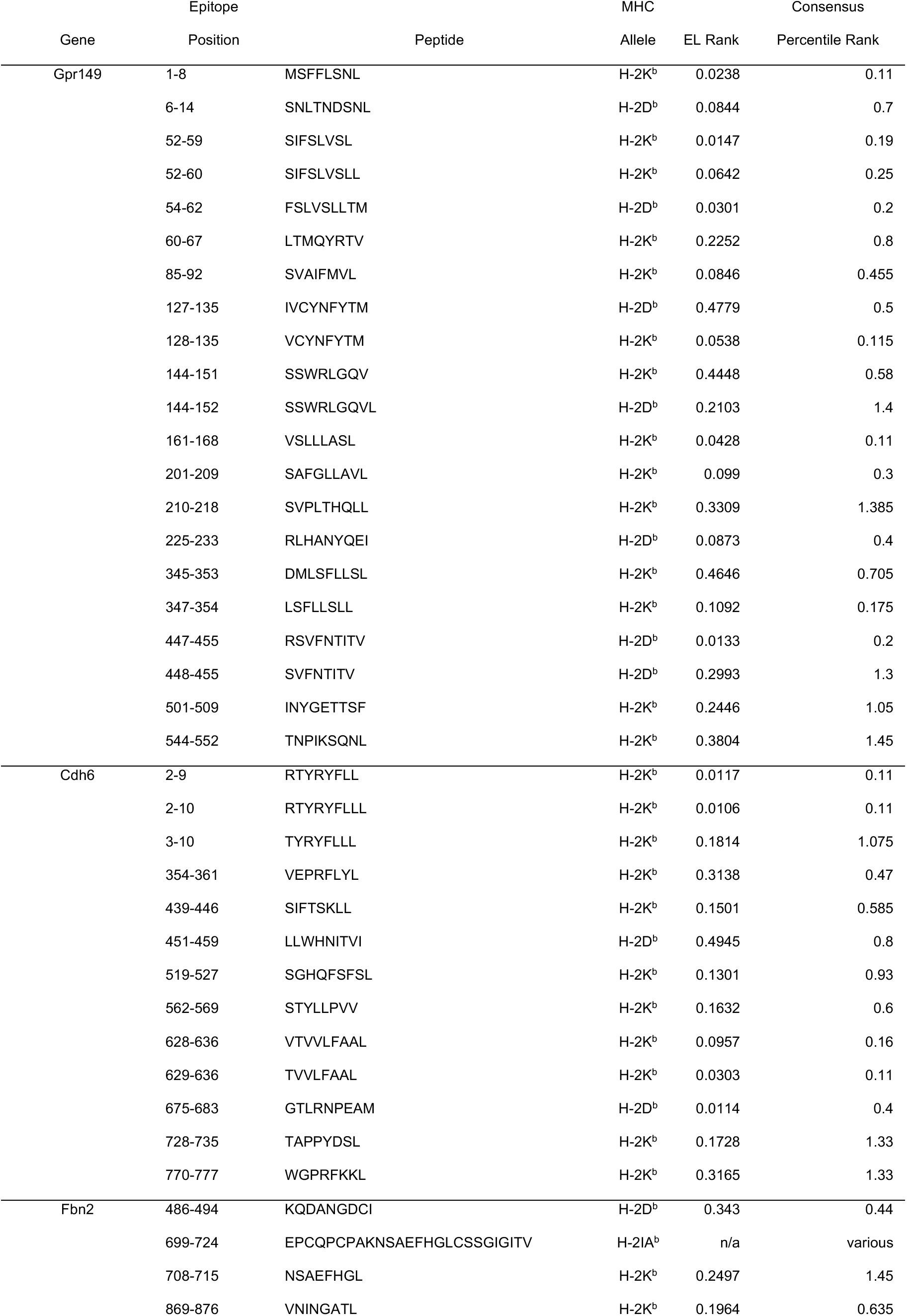
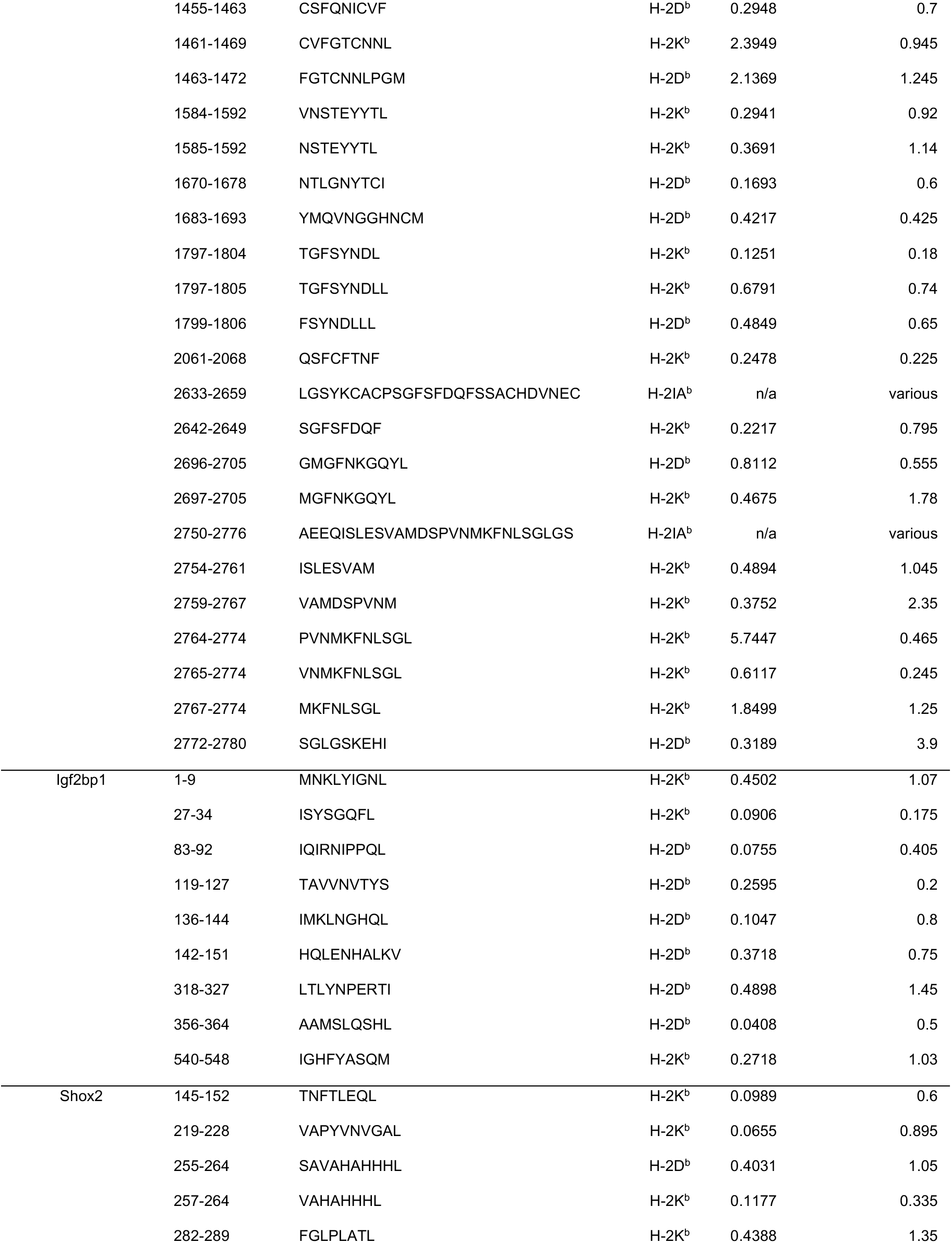
Peptides used for immunogenicity testing of TAAs. Amino acid position within original gene, peptide amino acid sequence, predicted MHC binding allele, and predicted MHC binding ranks are shown (EL Rank calculated by NetMHC4.1 Algorithm; Consensus Percentile Rank calculated by IEDB Consensus Algorithm). For Gpr149, Cdh6, Shox2, and Igf2bp1, all minimal epitopes with both an EL Rank < 0.5 and Consensus Percentile Rank < 1.5 were tested; for Fbn2, all predicted minimal epitopes within the minigene sequences with an EL Rank < 0.5 or Consensus Percentile Rank < 1.5 were tested. The top 48 H-2IA^b^ epitopes predicted by the IEDB MHC-II 2.22 Consensus Algorithm within the Fbn2 minigenes were all contained within three minigenes. Long 26- or 27-mer peptides corresponding to these three Fbn2 minigenes were also tested.

**Supplemental Table 7.**
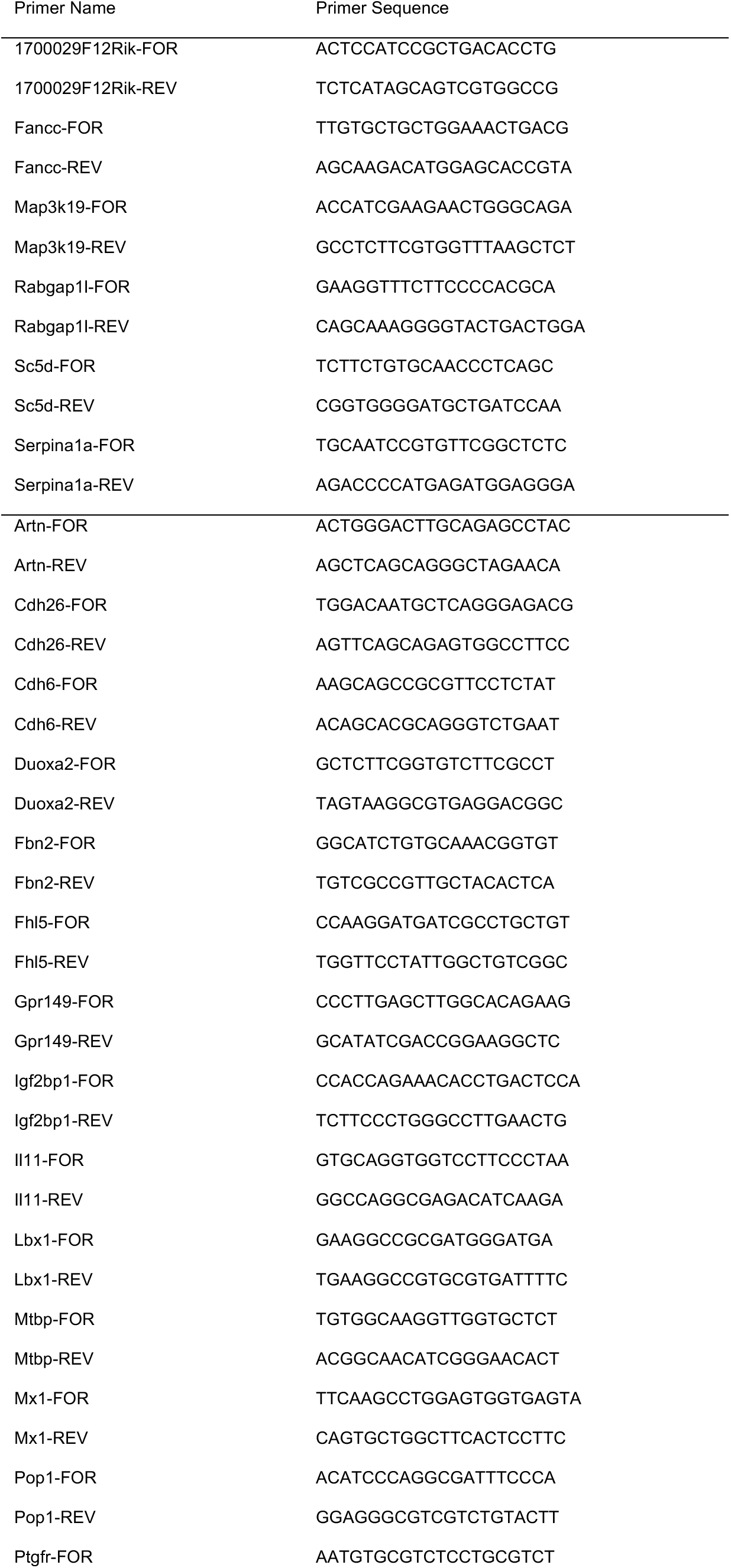
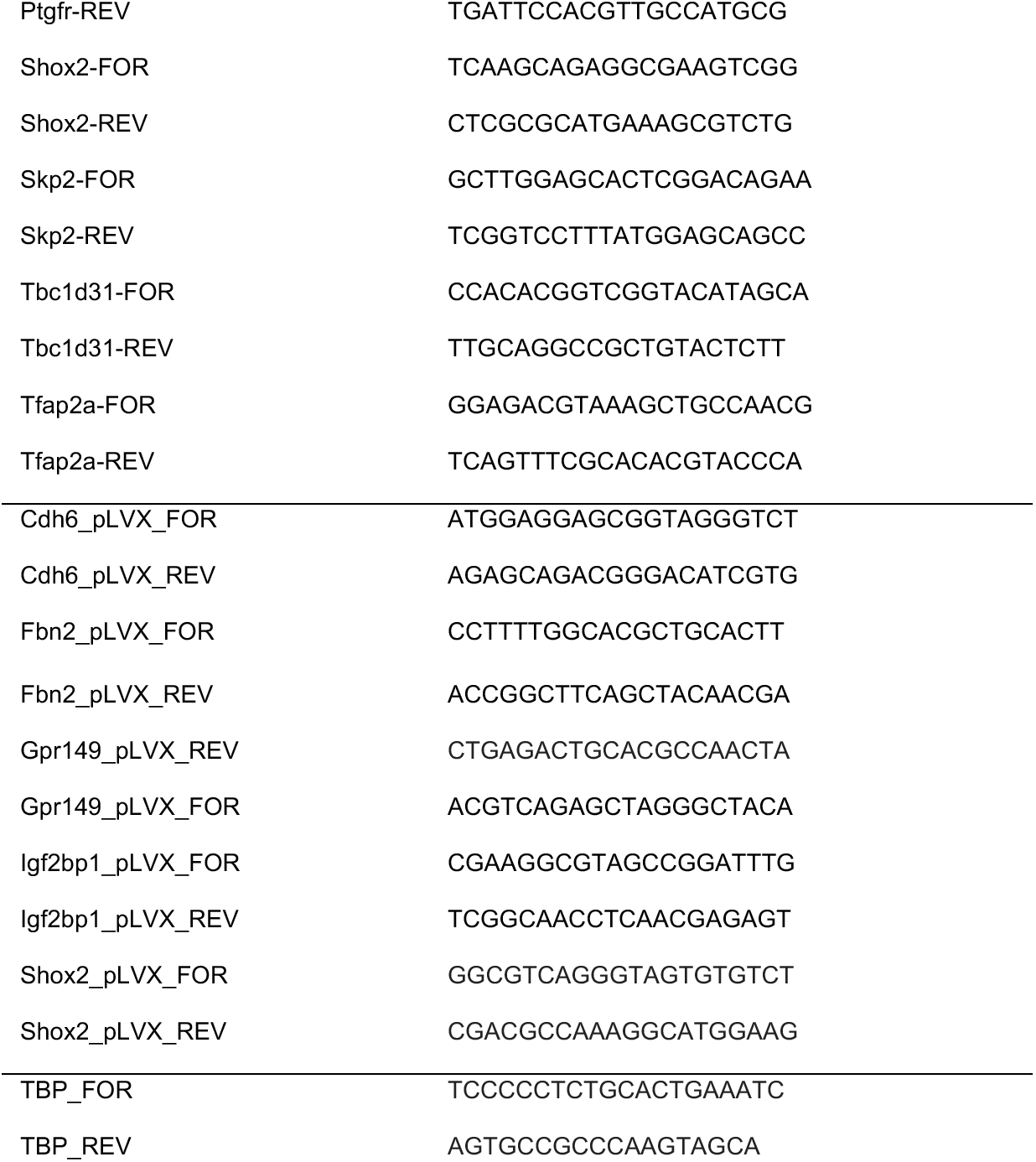
Primers used for RT-qPCR.

